# *Caulobacter* requires anionic sphingolipids and deactivation of *fur* to lose lipid A

**DOI:** 10.1101/2022.01.20.477143

**Authors:** Justin J. Zik, Sung Hwan Yoon, Ziqiang Guan, Gabriele Stankeviciute Skidmore, Ridhi R. Gudoor, Karen M. Davies, Adam M. Deutschbauer, David R. Goodlett, Eric A. Klein, Kathleen R. Ryan

## Abstract

Lipid A, the membrane-anchored portion of lipopolysaccharide, is an essential component of the outer membrane (OM) of nearly all Gram-negative bacteria. Here, we identify regulatory and structural factors that together permit *Caulobacter crescentus* to eliminate lipid A from its OM. Mutations in the ferric uptake regulator *fur* allow *Caulobacter* to survive in the absence of either LpxC, which catalyzes an early step of lipid A synthesis, or CtpA, a tyrosine phosphatase homolog which we find is needed for wild-type lipid A structure and abundance. Alterations in Fur-regulated processes, rather than iron status *per se*, underlie the ability to eliminate lipid A. Fitness of lipid A-deficient *Caulobacter* requires a previously uncharacterized anionic sphingolipid, ceramide phosphoglycerate (CPG), which also mediates sensitivity to the antibiotic colistin. Our results demonstrate that, in an altered regulatory landscape, anionic sphingolipids can support the integrity of a lipid A-deficient OM.

## Introduction

Gram-negative bacteria are enclosed in a three-layer envelope, composed of the inner or cytoplasmic membrane (IM), a thin layer of peptidoglycan (PG), and an outer membrane (OM). The OM is an asymmetric bilayer, with phospholipids populating the inner leaflet and lipopolysaccharide (LPS) predominating in the outer leaflet. The canonical LPS structure, first described in *Escherichia coli*, consists of three segments: 1) lipid A, a hexa-acylated, phosphorylated glucosamine disaccharide anchored in the membrane; 2) a core oligosaccharide usually shared by members of the same species; and 3) a repeating polysaccharide (O-antigen) which can vary highly among strains of the same species (Whitfield and Trent, 2014). LPS confers robust barrier function upon the OM, making it inherently less permeable than the IM to small hydrophobic compounds. Barrier function has been ascribed to strong lateral interactions between LPS molecules mediated by packing of the saturated acyl chains of lipid A and a network of divalent cations coordinated by negatively charged groups, particularly those on lipid A or the core polysaccharide (Nikaido, 2003).

Although the O-antigen and core polysaccharide are nearly always dispensable, it is widely accepted that the lipid A portion of LPS is essential for Gram-negative bacterial viability. Some exceptions to this rule are species that possess a dual membrane system but naturally lack lipid A, such as *Borrelia burgdorferi, Treponema pallidum*, and *Sphingomonas spp*. (Kawahara et al., 1991; Kawasaki et al., 1994; Radolf and Kumar, 2018). Efforts to eliminate lipid A from *E. coli* strains have demonstrated that the intermediate molecule lipid IV_A_ is sufficient for viability, but only if the strain also has compensatory mutations that promote the export of this species across the IM (Mamat et al., 2008; Meredith et al., 2006). To date lipid A-deficient mutants have been recovered in *Neisseria meningitidis*, *Moraxella catarrhalis*, and *Acinetobacter baumannii* (Moffatt et al., 2010; Peng et al., 2005; Steeghs et al., 1998). It remains unclear why at least a minimal lipid A structure is essential in some Gram-negative bacteria but not others.

Lipid A is synthesized by the highly conserved Raetz pathway (Whitfield and Trent, 2014), yet significant variation exists in lipid A structures. In many species, mechanisms exist to modify the 1- and 4L-phosphoryl groups of lipid A to decrease its negative charge and reduce susceptibility to cationic antimicrobial peptides (Moffatt et al., 2019). In a few species, replacement of the 1- and/or 4L-phosphoryl groups of lipid A with sugars is constitutive (De Castro et al., 2008; Plötz et al., 2000). The predominant lipid A species in the alphaproteobacterium *Caulobacter crescentus* (Smit et al., 2008) varies from that of *E. coli* (Qureshi et al., 1988) in that the central glucosamine disaccharide is replaced by two 2,3- diamino-2,3-dideoxy-D-glucopyranose (GlcN3N) residues, and the 1- and 4L-phosphates are replaced by galactopyranuronic acid (Gal*p*A) residues.

The tyrosine phosphatase homolog *ctpA* (for *<u>C</u>aulobacter* tyrosine <u>p</u>hosphatase <u>A</u>) is essential for viability and is implicated in cell envelope maintenance, but its molecular function remains unknown (Shapland et al., 2011). Depletion of *ctpA* causes extensive OM blebbing, failure to resolve PG at the division site, and cell death. Here, we show that *ctpA* is required for the wild-type structure and abundance of lipid A. A screen for suppressors of *ctpA* essentiality recovered strains with null mutations in the O-antigen biosynthetic pathway or in the ferric uptake regulator *fur*. Surprisingly, mutations in *fur* also permitted the deletion of *lpxC*, which encodes an otherwise essential enzyme catalyzing the first committed step in lipid A synthesis. Δ*ctpA* and Δ*lpxC* strains containing suppressor mutations have significantly reduced or undetectable levels of lipid A, respectively.

To identify mechanisms that promote survival in the absence of lipid A, we used random barcode transposon-site sequencing (RB-Tnseq) to identify genes important for fitness when lipid A synthesis is chemically inhibited. Interestingly, we obtained several hits in genes required for sphingolipid synthesis in *Caulobacter* (Stankeviciute et al., 2019, 2021). Since *Sphingomonads* naturally lack LPS and bear anionic sphingolipids on the cell surface (Kawasaki et al., 1994), we hypothesized that anionic sphingolipids could support viability in the *Caulobacter* Δ*lpxC* strain. Indeed, we identified a previously unknown sphingolipid species, ceramide phosphoglycerate, which is a critical fitness factor in the absence of lipid A. Further, we found that ceramide phosphoglycerate, rather than LPS, underlies *Caulobacter*’s sensitivity to the cationic antimicrobial peptide colistin.

## Results

### Suppressor mutations affecting fur or O-antigen synthesis permit the loss of ctpA

We used a CtpA depletion strain (see Methods, CtpA depletion strain) to identify mutations that would support *Caulobacter* viability in the absence of CtpA (Shapland et al., 2011). When depleting CtpA by transferring cells from xylose-supplemented rich medium (PYEX) to dextrose-supplemented rich medium (PYED), KR3906 exhibits division defects, significant OM blebbing, and death (**Fig. 1**). We UV-mutagenized KR3906, plated survivors on solid PYED medium to deplete CtpA, and grew the recovered colonies in PYED liquid medium to allow loss of the plasmid bearing *ctpA*. Isolates that had successfully lost the plasmid were identified via chloramphenicol sensitivity and the inability to PCR-amplify *ctpA* (**Fig. 1A**). Genome resequencing of 17 confirmed suppressors yielded 15 strains with mutations in nine genes predicted to participate in O-antigen biosynthesis, one strain with a single mutation in *fur*, encoding the ferric uptake regulator, and one strain harboring a mutation in *fur* along with a mutation in an O-antigen biosynthetic gene (**Table S1**). Due to the frequent occurrence of frameshift or nonsense mutations, we assumed that each mutation disrupted the function of the affected gene.

**Figure 1.**
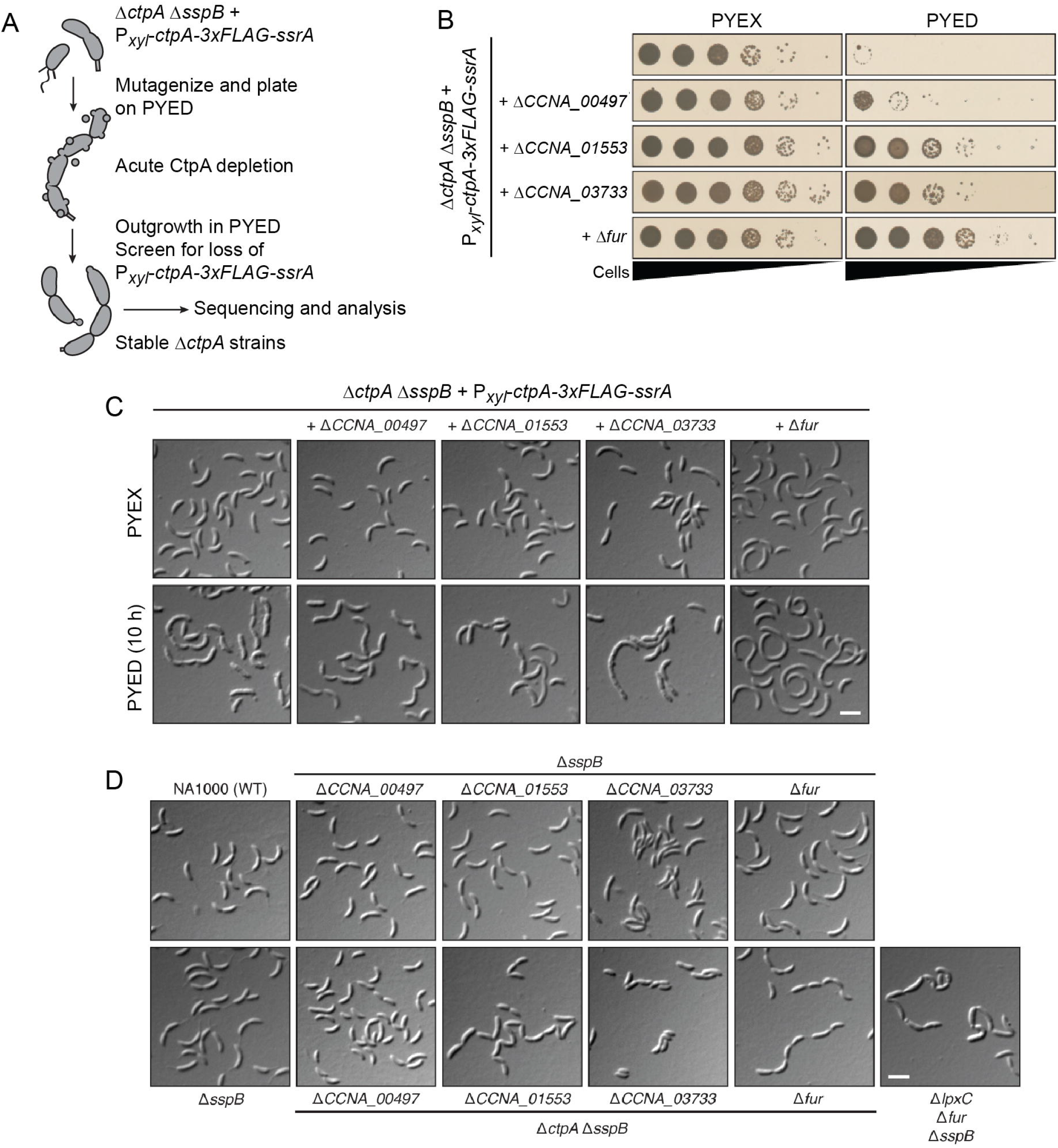
Suppressor mutations affecting *fur* or O-antigen biosynthesis permit the loss of *ctpA.* (A) Strategy for isolation of Δ*ctpA* suppressor mutants. (B) Viability of suppressor strains during CtpA depletion on PYED. (C) Differential interference contrast (DIC) images of the indicated strains grown in CtpA-expressing (PYEX) or non-expressing (PYED) conditions. D) DIC images of the indicated strains grown to exponential phase in PYE. Scale bar, 3 µm.

We chose the following candidate suppressors for further analysis: (i) *CCNA_00497*, encoding a predicted rhamnosyl transferase necessary for wild-type levels of O-antigen containing smooth LPS (S-LPS) (Hershey et al., 2019); (ii) *CCNA_01553*, encoding a glycosyltransferase which initiates O-antigen synthesis on undecaprenyl-phosphate (Toh et al., 2008); (iii) *CCNA_03733*, encoding a homolog of *manC* involved in synthesizing the activated sugar GDP-D-mannose (Samuel and Reeves, 2003), which is present in the core oligosaccharide and O-antigen of *Caulobacter* S-LPS (Jones et al., 2015); and iv) *CCNA_00055*, encoding the iron-responsive transcriptional regulator Fur (da Silva Neto et al., 2013). We individually deleted these genes in the CtpA depletion strain KR3906 while propagating the strains on PYEX.

To determine how each deletion affects cells during acute CtpA depletion, we shifted each mutant onto liquid or solid PYED medium and observed cell morphology and viability (**Fig. 1**). Compared to CtpA depletion in KR3906, acute depletion of CtpA in the Δ*fur* mutant caused much less OM blebbing, but still yielded elongated cells indicative of a division defect (**Fig. 1C**). Surprisingly, neither OM blebbing nor cell chaining/elongation was markedly improved when CtpA was depleted from the strains lacking *CCNA_00497*, *CCNA_01553*, or *CCNA_03733*. Despite the persistence of one or more morphological defects, deletion of *fur*, *CCNA_01553*, or *CCNA_03733* significantly improved cell viability during depletion of CtpA on solid PYED medium (**Fig. 1B**). In contrast, the deletion of *CCNA_00497* only weakly improved survival on PYED medium. Two independent point mutations in *CCNA_00497* were isolated in the suppressor screen, but each mutant also harbored 1-2 other mutations (**Table S1**) that may have contributed to the fitness of the original isolates.

To acquire stable Δ*ctpA* derivatives of each suppressor strain constructed above, we plated each new mutant on PYED medium and screened viable colonies for loss of the *ctpA*-bearing plasmid. The OM of each stable Δ*ctpA* strain was smooth with minimal blebbing, but chains of cells were still prevalent in the Δ*CCNA_01553* and Δ*fur* mutants (**Fig. 1D**). These reconstituted suppressor strains are morphologically similar to the original isolates containing point mutations in the same genes (**Fig. S1A**). Suppressed Δ than the wild-type strain NA1000 and the corresponding *ctpA^+^* strains, but all achieve similar stationary phase densities in PYE medium (**Fig. S1B**). As expected, restoring the expression of *fur*, *CCNA_00497*, or *CCNA_03733* using a xylose-inducible promoter reduced the viability of each corresponding stable Δ*ctpA* strain (**Fig. S1C**). Thus, null mutations affecting *fur* or O-antigen biosynthesis allow *Caulobacter* to survive in the absence of *ctpA*.

To confirm the functions of suppressor genes predicted to be involved in O-antigen synthesis, we deleted the individual genes in a control strain lacking *sspB* (see Methods, CtpA depletion strain). Whole-cell lysates treated with proteinase K were probed with antibodies recognizing S-LPS (**Fig. S2A**) or stained with Pro-Q Emerald 300 to detect carbohydrates (**Fig. S2B**). As previously observed, a control strain Δ*CCNA_01068* (*wbqA*) and the Δ*CCNA_01553* mutant lacked S-LPS (Awram and Smit, 2001), which migrates as a single high-molecular weight species in *Caulobacter* (Walker et al., 1994). Mutations in *CCNA_00497* or *CCNA_03733* caused partial or complete elimination of S-LPS, respectively. S-LPS was restored to each mutant by xylose-driven complementation of the respective genes (**Fig. S2C**). In contrast to strains with mutations in *CCNA_00497*, *CCNA_01553*, or *CCNA_03733*, the Δ*fur* Δ*sspB* mutant contained wild-type levels of S-LPS (**Fig. S2A-C**), indicating that *fur* mutations do not suppress the lethality of Δ*ctpA* by eliminating the O-antigen.

We propose that a low-molecular weight band in most lysates represents lipid A + core polysaccharide (**Fig. S2B and S2C, \*\***). This inference is supported by the band pattern in the Δ*CCNA_03733* (*manC*) mutant. The core oligosaccharide of *Caulobacter* LPS contains a single penultimate mannose residue (Jones et al., 2015); thus, the reduced size of the indicated band Δ*CCNA_03733* (**Fig. S2B and S2C, \***) may arise from an incomplete core oligosaccharide.

### Suppressor mutations permit ΔctpA and ΔlpxC strains to survive with little or no lipid A

*ctpA* is transcribed divergently from an operon containing *msbA*, *lpxJ*, *kdtA*, and *lpxK* (Zhou et al., 2015), which in other bacteria participate in the synthesis and export of lipid A + core polysaccharide (Whitfield and Trent, 2014). Like *ctpA*, these genes are essential for *Caulobacter* viability (Christen et al., 2011). Since CtpA depletion results in OM defects, and suppressor mutations were identified in O-antigen biosynthetic genes, we hypothesized that *ctpA* is required for some aspect of LPS synthesis or export.

We performed hot aqueous-phenol extraction of LPS from suppressor mutants lacking *ctpA*, along with their *ctpA^+^* counterparts, and analyzed them by PAGE and Pro-Q Emerald 300 staining. Full-length S-LPS was recovered from NA1000, Δ*sspB*, and Δ*fur* Δ*sspB* (**Fig. 2B, \*\*\***), whereas only smaller LPS species were present in the Δ*sspB* strains lacking *CCNA_00497*, *CCNA_01553*, or *CCNA_03733*. Interestingly, all Δ*ctpA* strains harboring suppressor mutations were deficient in low-molecular weight species that could represent lipid A +/- core oligosaccharide (**Fig. 2B**). We therefore used the Limulus Amebocyte Lysate (LAL) assay to measure lipid A abundance in live *Caulobacter* strains. The stable Δ approximately 1,000-fold less lipid A than NA1000 or the corresponding *ctpA^+^* strains (**Fig. 2A**).

**Figure 2.**
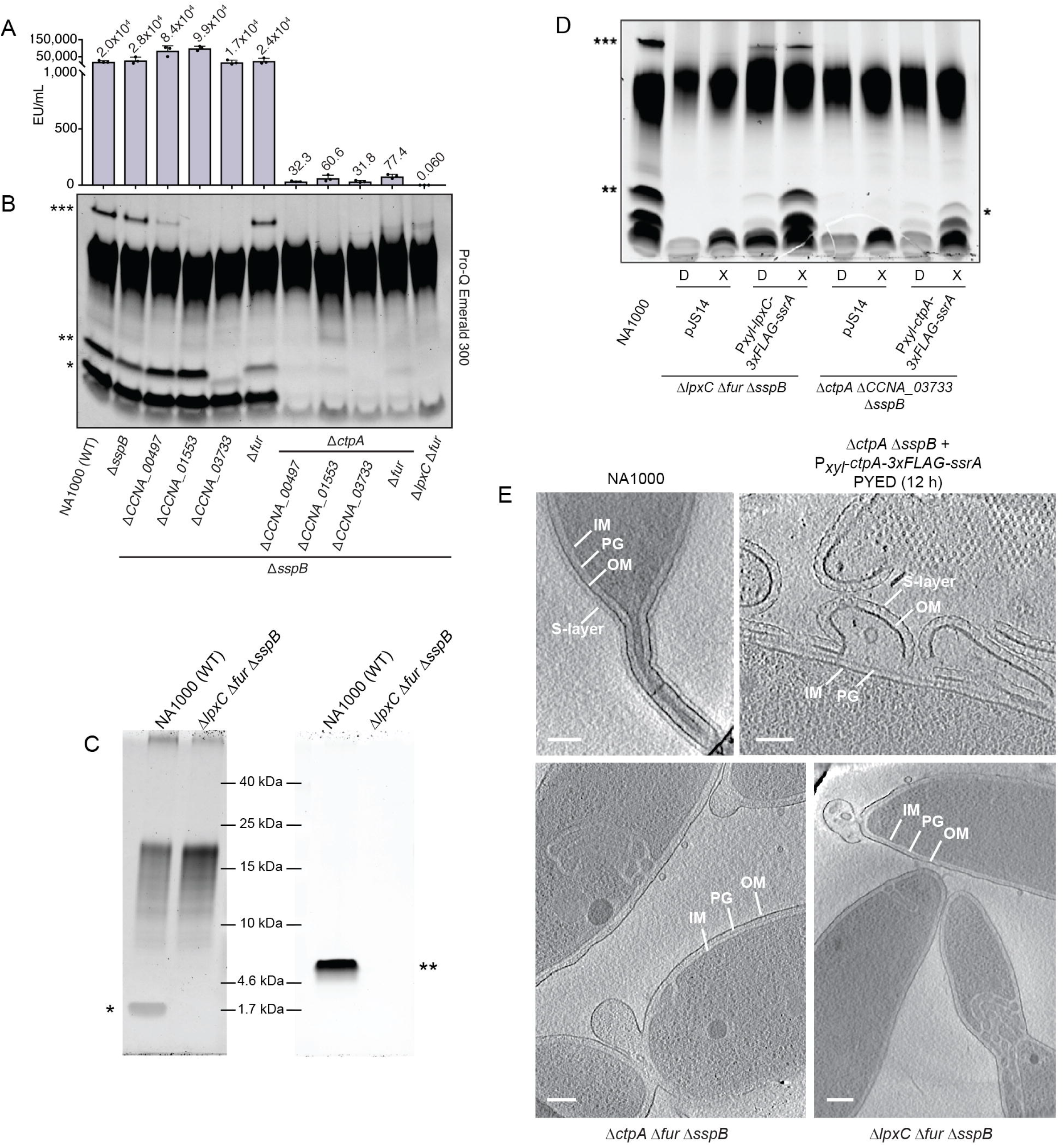
Δ*ctpA* and Δ*lpxC* strains with suppressor mutations contain little or no lipid A. (A) Endotoxin units (EU) per mL of whole cells of the strains indicated in panel B (mean ± S.D., *N*=3). Dots represent individual data points, and mean values are displayed above bars. (B) Hot aqueous-phenol LPS extracts of the indicated strains. *** = S-LPS, ** = putative full-length lipid A-core, * = putative lipid A. (C) Lipid A (left) or rough LPS (right) extracted from the indicated strains. * = lipid A. ** = rough LPS. (D) Proteinase K-treated lysates of the indicated strains. Cells were maintained in PYED or shifted into PYEX for 6 hours before harvesting. Samples were normalized by OD_660_. NA1000 was grown in PYE. Leaky expression of LpxC can generate S-LPS (***) and lipid A + core (**) in PYED. Full-length S-LPS is not restored to *ctpA* and the lipid A-core species is reduced in size (*) because *CCNA_03733* is needed for mannose incorporation. (E) Electron cryotomography images of the indicated strains noting the inner membrane (IM), peptidoglycan (PG), outer membrane (OM), and S-layer. All strains were grown to exponential phase in PYE medium, except that CtpA was depleted from KR3906 during 12 hours of growth in PYED prior to analysis. Scale bars, 100 nm.

Since Δ*ctpA* suppressor mutants could survive with drastically reduced amounts of lipid A, we asked if mutations in *fur* or O-antigen synthesis could allow *Caulobacter* to lose lipid A entirely. LpxC catalyzes the first committed step in lipid A synthesis, removal of the 2-acetyl group from acylated UDP-GlcNAc (Whitfield and Trent, 2014). The *lpxC* homolog *CCNA_02064* is essential for viability in wild-type *Caulobacter* (Christen et al., 2011). We constructed an LpxC depletion strain (KR4007) analogous to the CtpA depletion strain. We subsequently deleted *fur*, *CCNA_00497*, *CCNA_01553*, or *CCNA_03733* in this strain and examined the effects of acute LpxC depletion during growth in PYED. In the absence of candidate suppressor mutations, LpxC depletion yielded chains of cells with extensive membrane blebs. Cells lacking a gene for O-antigen synthesis still showed membrane blebs and chaining when LpxC was depleted (**Fig. 3A**). Cells lacking *fur* had far fewer OM blebs upon LpxC depletion but were still frequently elongated or chained (**Fig. 3A**). These morphologies are generally similar to those seen during CtpA depletion, but unlike CtpA, only Δ*fur* allowed significant growth of the LpxC depletion strain on solid PYED medium (**Fig. 3B**).

**Figure 3.**
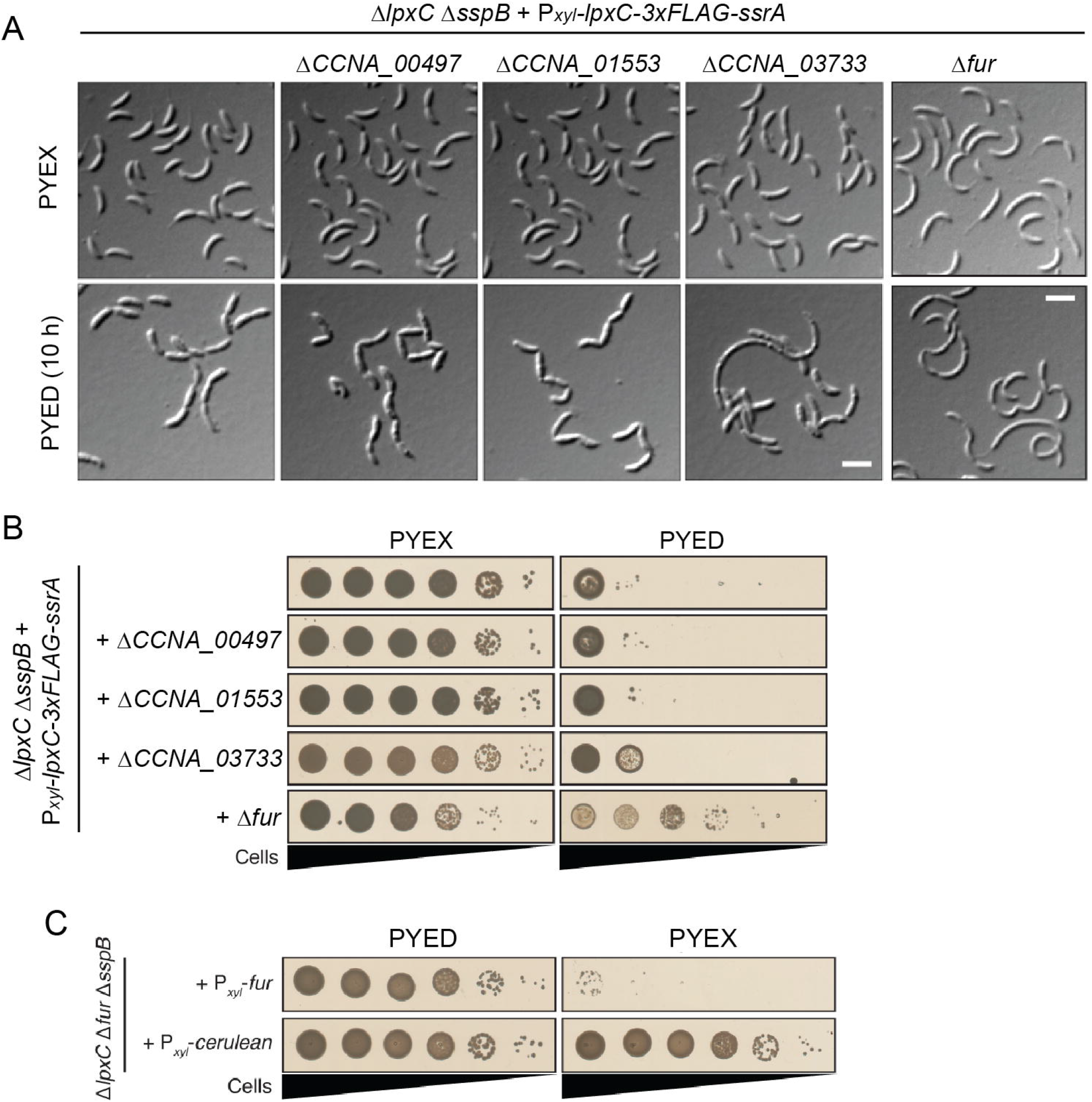
Deletion of *fur* supports the viability of Δ*lpxC* cells. (A) DIC images of the LpxC depletion strain alone or harboring the indicated mutations, grown in PYEX or PYED for 10 hours. Scale bar, 3 μm. (B) Viability of the LpxC depletion strain, alone or harboring the indicated mutations, plated on PYEX or PYED. (C) Viability of Δ*lpxC* Δ*fur* Δ*sspB* cells harboring a Pxyl-*fur* or a Pxyl-*cerulean* expression vector. Plates included kanamycin to retain expression vectors.

When we attempted to isolate stable Δ*lpxC* mutants using the same outgrowth and screening process as for Δ*ctpA*, only the strain harboring a Δ*fur* mutation permitted complete loss of *lpxC*. We initially recovered two stable Δ*lpxC* isolates from a depletion strain harboring Δ*CCNA_00497*, but genome resequencing revealed that these strains had acquired additional mutations in *fur* (**Table S2**). As in Δ*ctpA* Δ*fur* Δ*sspB*, the stable Δ*lpxC* Δ*fur* Δ*sspB* mutant still formed chains (**Fig. 1D**), and xylose-driven *fur* expression induced lethality in this background (**Fig. 3C**).

Background levels of lipid A were detected in Δ*lpxC* Δ*fur* Δ*sspB* cells in the LAL assay (**Fig. 2A**), strongly suggesting that lipid A is absent. To corroborate this result, we extracted LPS species by three distinct methods, separated them by PAGE, and stained with Pro-Q Emerald 300. Hot aqueous-phenol extracts of Δ*lpxC* Δ*fur* Δ*sspB* cells were deficient in S-LPS and putative lipid A +/- core species (**Fig. 2B**). However, unknown carbohydrate-containing species were extracted by this method. Extraction of free lipid A (El Hamidi et al., 2005) revealed that a species of ∼1800 Da, consistent with the mass of *Caulobacter* lipid A (Smit et al., 2008), is present in NA1000 but absent from Δ*lpxC* Δ*fur* Δ*sspB* (**Fig. 2C, left**). Again, however, unidentified carbohydrate species were present in these extracts. Lastly, the method of Darveau and Hancock (Darveau and Hancock, 1983) yielded a single rough LPS species which was present in NA1000 and absent from Δ*lpxC* Δ*fur* Δ*sspB* (**Fig. 2C, right**); this method resulted in no unidentified contaminants. Although some *Caulobacter* extracts contain unidentified lipids, these assays together strongly imply that lipid A is absent from the Δ*lpxC* Δ*fur* Δ*sspB* mutant. Xylose-driven expression of *lpxC* or *ctpA* restored the production of lipid A-containing species to Δ*lpxC* Δ*fur* Δ*sspB* or Δ*ctpA* Δ*CCNA_01553* Δ*sspB*, respectively (**Fig. 2D**).

Lipid A extracts from Δ*ctpA* Δ*fur* Δ*sspB*, Δ*lpxC* Δ*fur* Δ*sspB*, a nd control strains were further analyzed by matrix-assisted laser desorption/ionization tandem mass spectrometry (MALDI-MS/MS). Wild-type NA1000, Δ*sspB*, and Δ*fur* Δ*sspB* extracts contained predominantly the full-length lipid A (*m/z* 1874, (Smit et al., 2008) and lesser amounts of an ion at *m/z* 1858 that differs from 1874 by 16 *m/z*, consistent with the absence of one hydroxyl group (**Fig. S3A- C**). MALDI-MS analyses of lipid A extracts from Δ*ctpA* Δ*fur* Δ*sspB* cells revealed no ions consistent with full-length *Caulobacter* lipid A, but identified ions at *m/z* 1682 and *m/z* 1486 (**Fig. S3D**) that appeared to be missing the Gal*p*A residues at the 1 and 4L positions. Tandem MS analysis of these ions revealed mass losses consistent with phosphates, as would be expected for canonical lipid A structures. Although additional characterization is needed, our results suggest that *Caulobacter* mutants lacking CtpA produce an incomplete lipid A species which retains phosphate at the 1 and 4L positions, and which lacks one or more of the secondary phospholipids. While these incomplete lipid A species were detectable by mass spectrometry, gel electrophoresis and LAL assays indicate that they are much less abundant than the lipid A in wild-type strains.

Lipid A extracts from the Δ*lpxC* Δ*fur* Δ*sspB* mutant yielded no ions consistent with wild-type lipid A and instead contained an unknown lipid (**Fig. S3E**, *m/z* 1412). Numerous attempts to interpret the structure of this ion using the same type of tandem MS data as used in Fig. S3A-D failed to generate a structural hypothesis resembling obvious choices such lipid A derivatives or cardiolipin. Again, it is important to note that while this unknown ion was detected by mass spectrometry, gel electrophoresis and LAL assays together indicate that lipid A is absent from Δ*lpxC* Δ*fur* Δ*sspB* cells.

### Lipid A-deficient Caulobacter mutants produce a three-layer cell envelope

We analyzed NA1000, Δ*ctpA* Δ*fur* Δ*sspB* and Δ*lpxC* Δ*fur* Δ*sspB* strains via electron cryotomography to assess the effects of mutations on cell envelope structure (**Movies S1-S4**). As expected, the S-layer is absent from both mutants due to the loss of its O-antigen attachment site (Walker et al., 1994). Despite drastic reductions in lipid A levels, the Δ*ctpA* Δ*fur*Δ*sspB* and Δ*lpxC* Δ*fur* Δ*sspB* mutants still generate a three-layer cell envelope, including an OM (**Fig. 2E**). Although much less severe than during acute CtpA depletion (**Movie S4**), membrane blebs were often observed at the cell poles or division sites in Δ*ctpA* Δ*fur* Δ*sspB* and membrane blebs were often observed at the cell poles or division sites in Δ*ctpA* Δ*fur* Δ*sspB* and exhibited defects in stalk structure or IM distortions at the pole or midcell (N = 100; Δ*ctpA* Δ*fur* exhibited defects in stalk structure or IM distortions at the pole or midcell (N = 100; Δ*ctpA* Δ*fur*Δ*sspB*: 61%; Δ*lpxC* Δ*fur* Δ*sspB*: 51%; NA1000: 4%).

### Fur-regulated processes, rather than available iron levels, control the conditional essentiality of lipid A

LPS defects are usually associated with increased chemical sensitivity (Nikaido, 2003). Mutations in *fur* or O-antigen synthesis genes did not appreciably increase chemical sensitivity compared to NA1000, while strains lacking *ctpA* or *lpxC* had greater sensitivity to a subset of antibiotics and to all tested detergents (**Fig. 4A**). In sharp contrast, the Δ*lpxC* Δ*fur* Δ*sspB* strain and Δ*ctpA* Δ*sspB* strains with suppressor mutations were much less susceptible to CHIR-090, an inhibitor of LpxC (McClerren et al., 2005) (**Fig. 4B**). We infer that suppressed Δ*lpxC* and Δ*ctpA* mutants are relatively insensitive to CHIR-090 because they already produce little lipid A or lack the target enzyme.

**Figure 4.**
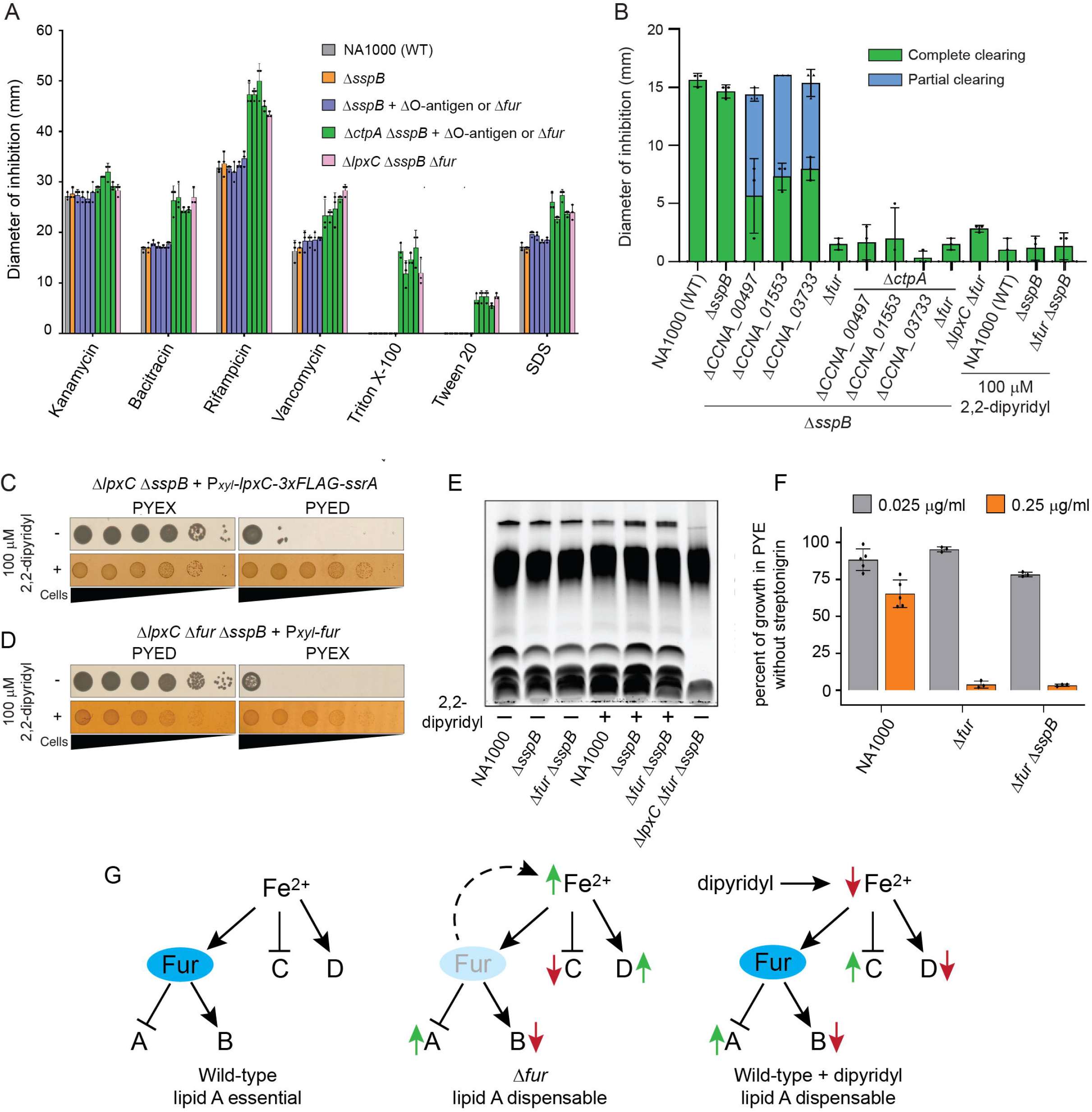
Fur-regulated processes control the conditional essentiality of lipid A. (A) Sensitivity to the indicated chemicals measured by disc diffusion assay. Dots indicate individual data points. Suppressor mutations present in strains represented by blue or green bars are, from left to right, Δ*CCNA_00497*, Δ*CCNA_01553*, Δ*CCNA_03733*, and Δ*fur*. (B) CHIR-090 sensitivity measured by disc diffusion assay. Partial clearing indicates the diameter of a ring of intermediate growth. Dots and triangles indicate individual measurements of cleared and partially cleared zones, respectively (mean ± S.D., N=3). (C) Viability of the LpxC depletion strain in inducing (PYEX) or depleting (PYED) conditions, in the presence or absence of 100 µM 2,2’-dipyridyl. (D) Viability of the Δ*lpxC* Δ*fur* Δ*sspB* strain harboring a Pxyl-*fur* plasmid, grown in noninducing (PYED) or inducing (PYEX) conditions, in the presence or absence of 100 µM 2,2’-dipyridyl. Plates included kanamycin to retain the expression vector. Brightness was reduced and contrast increased to improve the clarity of colonies grown on 2,2’-dipyridyl. (E) Proteinase K-treated lysates of the indicated strains grown overnight in the presence or absence of 100 μ dipyridyl. Samples were normalized by OD_660_. (F) Growth inhibition by SNG in liquid PYE cultures of the indicated strains. Dots represent individual OD_660_ ratios (mean ± S.D.). (G) Genes regulated by Fur in concert with iron (sets A and B) are modulated similarly by deletion of *fur* or by iron limitation, while genes regulated by iron alone (sets C and D) are modulated in opposite directions. Changes in Fur-regulated gene expression correlate with the ability to lose lipid A.

In agreement with its ability to suppress the lethality of Δ*lpxC* and Δ*ctpA* mutations, the Δ*fur* allele by itself greatly reduced the sensitivity of *Caulobacter* to CHIR-090 (**Fig. 4B**). Fur senses available Fe^2+^ by reversibly binding a [2Fe-2S] cluster (Fontenot et al., 2020). We therefore asked whether iron limitation could mimic the phenotypes of a Δ*fur* mutant. Culturing NA1000 with the iron chelator 2,2’-dipyridyl reduced its susceptibility to CHIR-090 to match that of the Δ*fur* mutant (**Fig. 4B**). Neither depleting LpxC in *fur^+^* cells nor inducing *fur* in Δ*lpxC* Δ*fur* Δ*sspB* cells caused a reduction in viability in the presence of 2,2’-dipyridyl (**Fig. 4C, D**). The NA1000, Δ*sspB*, and Δ*fur* Δ*sspB* strains cultured in 2,2′-dipyridyl retained LPS and lipid A +/- core species (**Fig. 4E**). Therefore, low iron availability does not induce the loss of lipid A, but is sufficient to maintain *Caulobacter* viability when lipid A is eliminated by chemical or genetic means.

In diverse bacteria, Fur inhibits the expression of iron uptake systems and promotes the expression of proteins that utilize iron, contributing to iron homeostasis (Andrews et al., 2013). Because they are impaired in iron sensing, *fur* mutants of other bacteria accumulate more available iron than the corresponding wild-type strains (Liu et al., 2020; Wofford et al., 2019). We measured available iron levels using a streptonigrin (SNG) sensitivity assay (Justino et al., 2007; Nachin et al., 2001), because SNG killing is linked to the intracellular formation of oxygen radicals in the presence of iron (Hassett et al., 1987; Yeowell and White, 1982). Growth of the Δ*fur* and Δ*fur* Δ*sspB* strains was almost completely inhibited by 0.25 μg/ml SNG, while NA1000 was only mildly inhibited (**Fig. 4F**), consistent with higher levels of available iron in Δ These findings indicate that both excess available iron (in *fur* mutants) and iron depletion (by 2,2-dipyridyl) are compatible with the elimination of lipid A. Since *fur* deletion and iron chelation have the same effect on Fur-regulated gene expression, but opposite effects on Fur-independent iron signaling (**Fig. 4G**), this implies that iron-mediated processes regulated by Fur are specifically responsible for the survival of lipid A-deficient *Caulobacter* (Leaden et al., 2018; da Silva Neto et al., 2013).

### RB-Tnseq identifies sphingolipid synthesis genes needed for fitness when lipid A synthesis is chemically inhibited

To identify additional factors that promote the survival of lipid A-deficient *Caulobacter*, we challenged an RB-Tnseq library constructed in NA1000 (Price et al., 2018) with CHIR-090. Individual barcode frequencies were measured by high-throughput sequencing before each trial and after growth in either PYE or PYE + 2 μg/ml CHIR-090. To identify genes that are particularly important when LpxC is inhibited, we averaged and compared the gene fitness scores (Wetmore et al., 2015) from three trials in each condition (**Fig. 5A**). We anticipated that mutations in *fur* would increase fitness in CHIR-090, but the NA1000 RB-Tnseq library contained no insertions in *fur*. Surprisingly, nearly all genes known to be regulated by Fur (Leaden et al., 2018; da Silva Neto et al., 2013) had similar fitness scores in unstressed and CHIR-090-exposed cultures (**Fig. 5A**).

**Figure 5.**
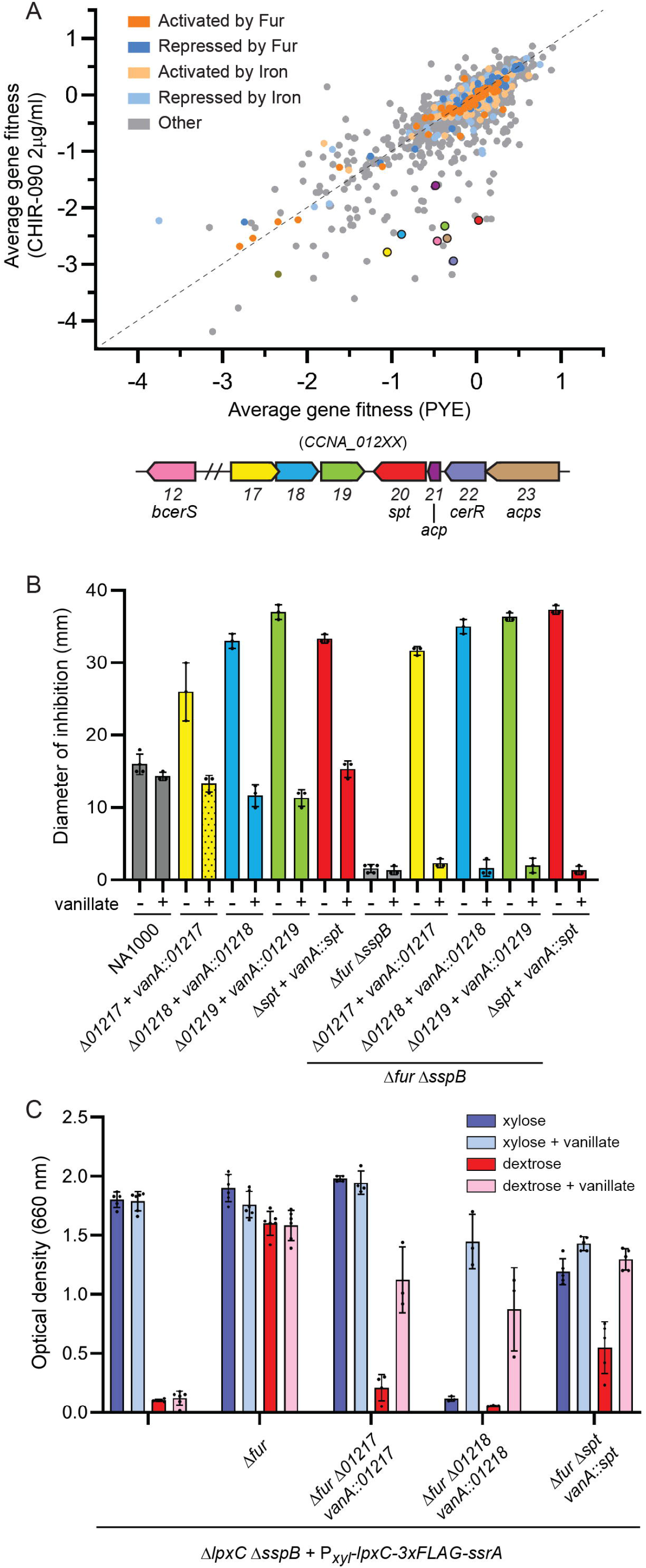
RB-Tnseq identifies sphingolipid synthesis genes needed for fitness when LpxC is inhibited. (A) Average gene fitness scores for three challenges of the NA1000 RB-Tnseq library with PYE or PYE+2 μg/ml CHIR-090. Fitness scores are color-coded based on regulation of the corresponding genes. Fitness scores of selected genes (not regulated by Fur or iron) are indicated by colors matching the open reading frame diagram below. (B) CHIR-090 sensitivity of the indicated strains measured by disc diffusion assay (mean ± S.D.) Where indicated, 0.5 mM vanillate was included in the medium. (C) Overnight growth of strains expressing (xylose) or depleting (dextrose) LpxC, and expressing (vanillate) or not expressing the indicated genes. Dots represent individual measurements (mean ± S.D.).

Focusing on genes whose average fitness scores were ≥ 1 point lower in CHIR-090-treated cultures than in control cultures, we identified five genes involved in sphingolipid synthesis: *spt* (*CCNA_01220*), acyl-carrier protein (*acp*, *CCNA_01221*), *cerR* (*CCNA_01222*), ACP-synthetase (*acps*, *CCNA_01223*), and *bcerS* (*CCNA_01212*) (Olea-Ozuna et al., 2021; Stankeviciute et al., 2021). Additionally, fitness scores were lower for transposon insertions in a neighboring operon of three uncharacterized genes predicted to modify lipids (*CCNA_01217- 01219*, Marks et al., 2010) (**Fig. 5A**). To examine the roles of genes in the uncharacterized operon, we constructed unmarked deletions in the NA1000 and Δ*fur* Δ*sspB* backgrounds and complemented them with the corresponding genes expressed from the inducible *vanA* promoter (Thanbichler et al., 2007). Loss of *spt*, *CCNA_01217*, *CCNA_01218*, or *CCNA_01219* greatly increased the susceptibility to CHIR-090, either in NA1000 or in Δ*fur* Δ*sspB* cells (**Fig. 5B**), and expression of the complementing gene from the *vanA* locus restored the wild-type level of susceptibility, validating the RB-Tnseq results.

Mutations in *CCNA_01217-01219* or *spt* could increase CHIR-090 sensitivity via distinct mechanisms: by damaging the cell’s permeability barrier and giving easier access to CHIR-090, by making it more difficult for cells to grow after lipid A synthesis is inhibited, or both. To eliminate changes in drug access as a factor in the experiment, we measured the effects of *CCNA_01217*, *CCNA_01218,* and *spt* upon cell viability when LpxC was depleted. We deleted individual genes in the strain Δ*lpxC* Δ*fur* Δ*sspB* + P*xyl*-*lpxC::3xFLAG-ssrA* (KR4091) and complemented them with *vanA*-driven copies as described above. The parent strain lacks *fur* and grows in PYED medium when LpxC is depleted. In contrast, KR4091 lacking *CCNA_01217* or *spt* grew poorly in PYED medium, and growth in PYED was fully or partially restored by expressing the complementing gene from the vanillate promoter (**Fig. 5C**). KR4091 lacking *CCNA_01218* grew poorly in PYEX and PYED media without vanillate, indicating that KR4091 requires *CCNA_01218* for fitness even when LpxC is produced. Growth was improved by vanillate-driven expression of *CCNA_01218* (**Fig. 5C**). Since this assay does not rely on an exogenous chemical, we conclude that *CCNA_01217-8* and *spt* are critical for the fitness of lipid A-deficient *Caulobacter*, not simply for the exclusion of CHIR-090.

### CCNA_01217-01219 convert neutral ceramide to an anionic sphingolipid, ceramide phosphoglycerate

The importance of Spt for viability in the absence of lipid A indicated a role for sphingolipids in this phenotype. Since *Sphingomonads* produce anionic glycosphingolipids (GSLs) on the outer membrane (Kawasaki et al., 1994), we initially hypothesized that *Caulobacter* responds to *lpxC* deletion by upregulating GSL production. However, neither *Caulobacter* sphingolipid glycosyltransferase (*sgt1* or *sgt2*, Stankeviciute et al., 2019) was important for the fitness of CHIR-090-treated cells. A careful analysis of the *Caulobacter* lipidome revealed a previously unidentified sphingolipid species, ceramide phosphoglycerate (**Fig. 6A**). In fact, we identified two forms of this lipid containing either one or two phosphoglycerate moieties (**Fig. 6A**) that we have designated CPG and CPG2. LC/MS/MS analysis confirmed the proposed structures of these lipids (**Fig. 6B**).

**Figure 6.**
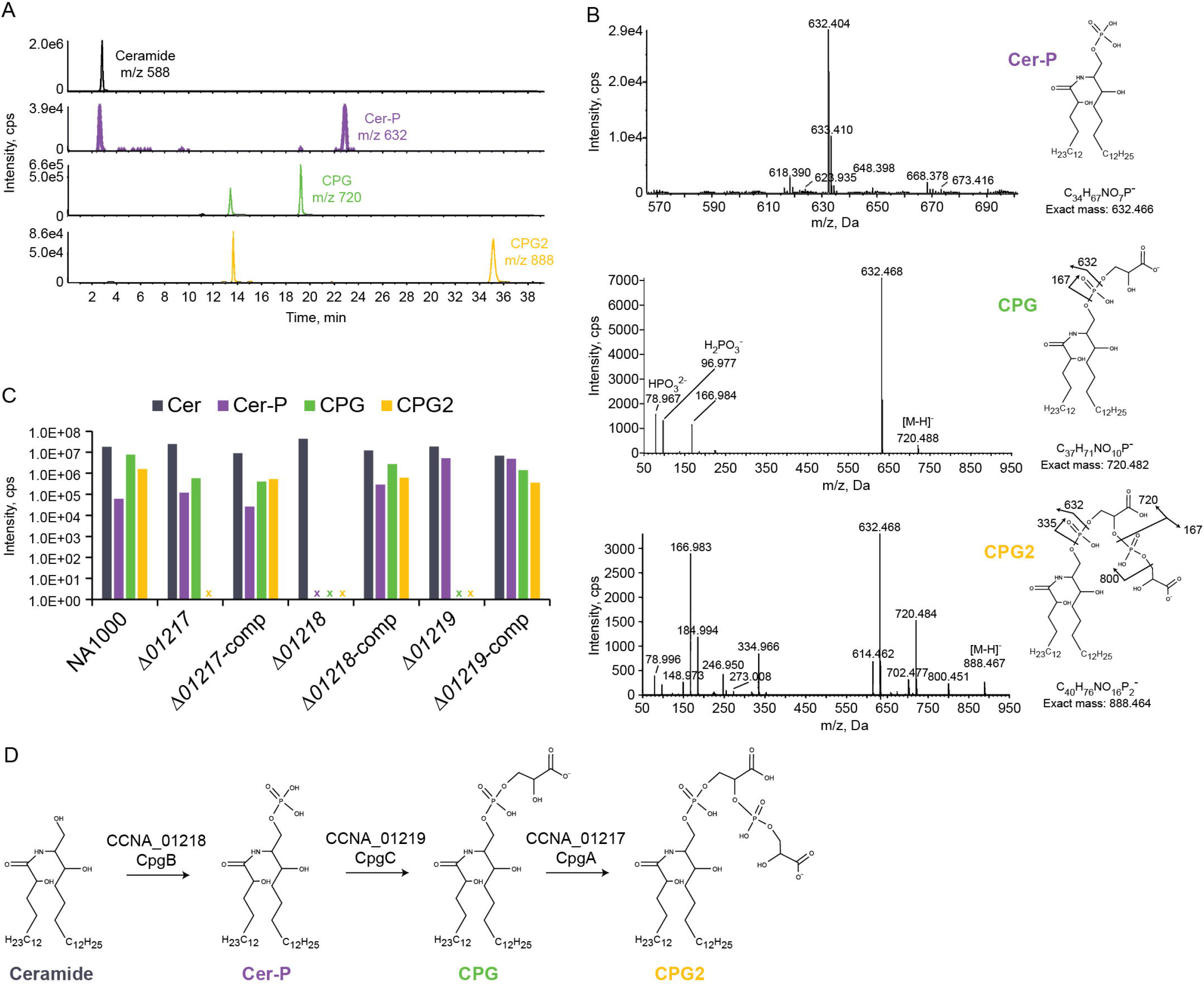
*CCNA_01217-01219* convert neutral ceramide to an anionic sphingolipid, ceramide phosphoglycerate. (A) Extracted ion-chromatograms identified the indicated sphingolipid species in NA1000 lipid extracts. (B) Structural determination of anionic sphingolipids was performed by MS/MS analysis. (C) The presence of the indicated sphingolipids was assessed in each deletion mutant and its respective complemented strain. x, no lipid of that type was detected in the indicated strain. (D) Proposed mechanism for CPG2 synthesis.

To determine whether *CCNA_01217-01219* are involved in CPG/CPG2 synthesis, we analyzed lipid extracts from mutant and complemented mutant strains. Deletion of *CCNA_01217* resulted in the loss of CPG2 but had no effect on CPG (**Fig. 6C**). CCNA_01217 has a conserved phosphatidylglycerophosphate synthase (PgsA) domain which is normally involved in phosphatidylglycerol (PG) synthesis. PG is the dominant phospholipid in *Caulobacter* (Stankeviciute et al., 2019), but the essential PgsA ortholog *CCNA_03002* is likely responsible for PG synthesis (Christen et al., 2011; Marks et al., 2010). Thus, we conclude that CCNA_01217 adds the second phosphoglycerate to form CPG2. Deletion of *CCNA_01218* led to the loss of CPG, CPG2 and ceramide-phosphate (**Fig. 6C**). CCNA_01218 is annotated as a sphingosine kinase-related protein and has a conserved LCB5 domain (Nagiec et al., 1998). Therefore, we propose that CCNA_01218 adds the initial phosphate on the ceramide. Lastly, the Δ*CCNA_01219* mutant lost CPG and CPG2 but retained ceramide-phosphate (**Fig. 6C**). This is consistent with CCNA_01219 adding a glycerate molecule to ceramide-phosphate to form CPG. CCNA_01219 has no conserved domains, and a BLAST analysis identified homologs only in Caulobacterales and Sphingomonodales. Each deletion could be complemented by expressing the respective gene from a vanillate-inducible promoter (**Fig. 6C**).

Based on these data, we propose a mechanism where CCNA_01218 (CpgB) phosphorylates ceramide, CCNA_01219 (CpgC) adds a glycerate, and CCNA_01217 (CpgA) adds a second phosphoglycerate (**Fig. 6D**). We note that the amount of CPG/CPG2 detected appears to be a relatively small percentage of the total lipidome (**Fig. S4**), raising the question of how these lipids can enable survival in the absence of lipid A. The CPG2 molecule is very polar, as evidenced by its very long LC retention time, and we expect that this lipid is not efficiently extracted by standard methods. Though we tried several modifications to increase the extraction yield, we made only marginal improvements. Additionally, our genetic data show that CpgA adds the second phosphoglycerate molecule to generate CPG2, but we cannot rule out the possibility of higher-order polymers containing additional phosphoglycerates, which would be even more polar and difficult to extract.

### Ceramide phosphoglycerate mediates susceptibility to colistin

Cationic antimicrobial peptides (CAMPs) have been demonstrated to kill Gram-negative bacteria by first interacting with negatively charged groups on surface-exposed LPS. Phosphates at the 1 and 4L positions of lipid A are particularly important for this interaction, and several bacteria possess mechanisms to modify them, reducing their negative surface charge and sensitivity to CAMPs (Moffatt et al., 2019; Velkov et al., 2010). Despite lacking phosphate groups on its lipid A, *Caulobacter* is highly sensitive to colistin (**Fig. 7A**) and the antimicrobial effect is retained in the lipid A-deficient strain Δ*lpxC* Δ*fur* Δ*sspB* (**Fig. 7A**). Since the CPG/CPG lipids are anionic, we considered whether they may be the colistin target in *Caulobacter*. Indeed, the growth of mutants lacking *cpgA*, *cpgB*, or *cpgC* was unaffected by colistin (**Fig. 7A**). Since the deletion of *cpgA*, catalyzing the conversion of CPG to CPG2, can alone greatly reduce colistin sensitivity, and since the elimination of lipid A had no effect, we infer that a primary target of colistin on the *Caulobacter* surface is CPG2. Furthermore, these findings are consistent with our hypothesis that CPG lipids are a significant component of the OM whose detection is limited by inefficient extraction.

**Figure 7.**
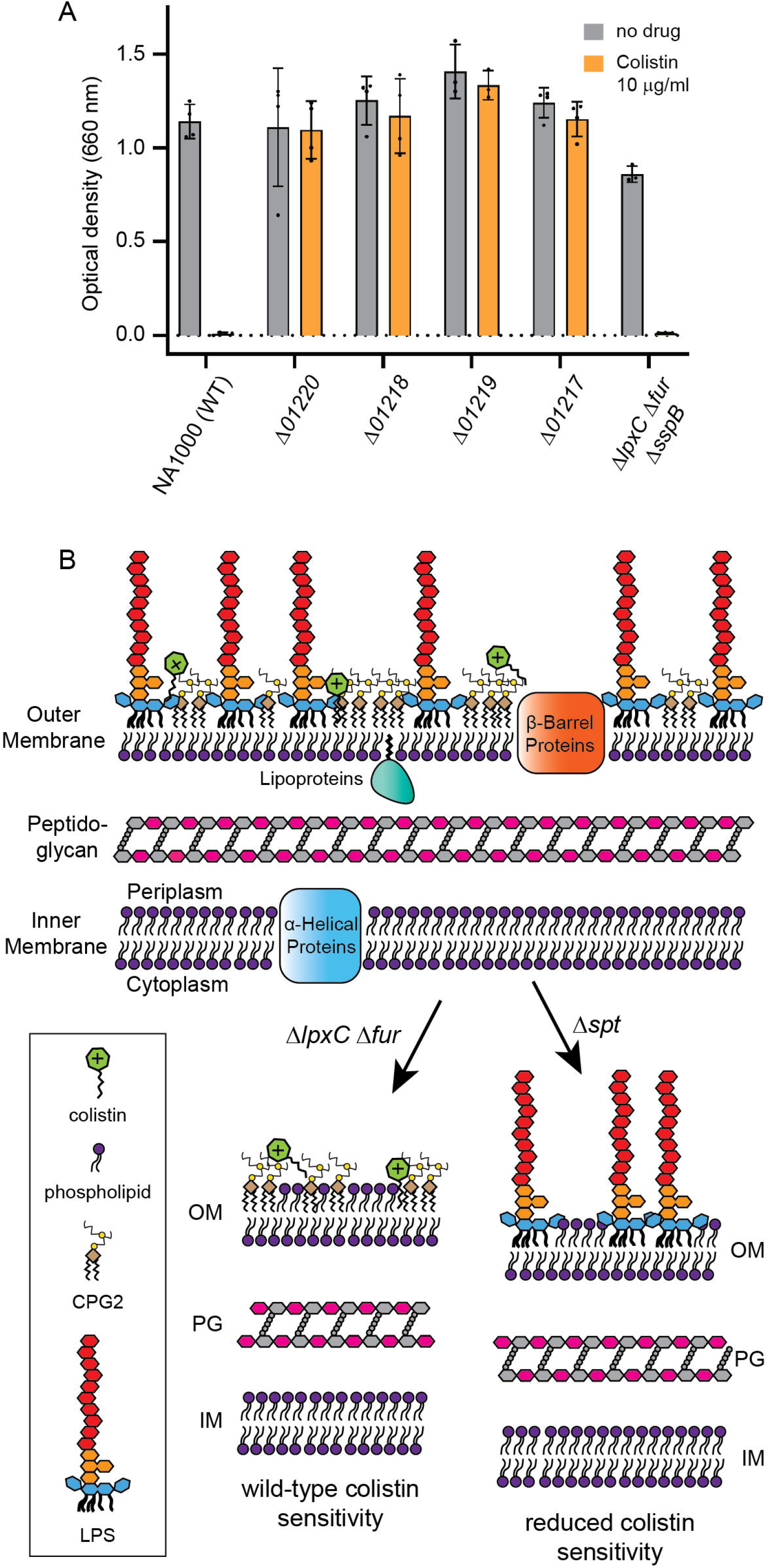
Ceramide phosphoglycerate mediates susceptibility to colistin. (A) Overnight growth of the indicated strains in the presence or absence of 10 μg/ml colistin. Dots represent individual OD_660_ measurements (mean ± S.D.). (B) Model of the *Caulobacter* cell envelope containing LPS and CPG2. Consequences for OM composition and colistin sensitivity when either lipid A (Δ*lpxC* Δ*fur*) or sphingolipids (Δ*spt*) are eliminated. The presence of CPG2 and the absence of *fur* are together required for the viability of lipid A-deficient *Caulobacter*.

## Discussion

### CtpA is required for wild-type lipid A structure and abundance

We performed a suppressor screen to discover the essential function of CtpA, which contains active site residues characteristic of tyrosine phosphatases. Inactivation of *fur* or genes involved in O-antigen synthesis permitted the deletion of *ctpA* and yielded cells with drastically reduced amounts of lipid A. MS/MS analysis of the remaining lipid A extracted from Δ*ctpA* Δ*fur* Δ*sspB* were consistent with species that retain phosphoryl groups at the 1 and 4L positions of the central disaccharide, suggesting that CtpA is responsible for dephosphorylating at least one of these positions, in preparation for the addition of Gal*p*A residues.

Some alphaproteobacteria produce lipid A species with a tri- or tetrasaccharide backbone (De Castro et al., 2008). In *Rhizobia*, the phosphatases LpxE and LpxF dephosphorylate the 1 and 4L positions, respectively, of lipid A at the periplasmic surface of the IM (Karbarz et al., 2003; Wang et al., 2006). Sugars are then added to the 1 and 4L positions by the glycosyltransferases RgtF and RgtD, respectively, before the transport of mature LPS molecules to the OM (Brown et al., 2012, 2013). NA1000 harbors a gene (*CCNA_03113*) with similarity to LpxE, but none with similarity to LpxF, raising the possibility that CtpA substitutes for LpxF. Additional work is required to verify this model; however, dephosphorylation of a lipid A intermediate by CtpA could explain the ability of mutations in O-antigen synthesis to suppress Δ -phosphates impairs lipid A trafficking, then removal or reduction of the O-antigen could partially compensate and allow at least a small amount of structurally altered lipid A to reach the cell surface.

### The elimination of lipid A in Caulobacter crescentus requires a novel anionic sphingolipid

Here we demonstrate that the enzyme LpxC and lipid A itself are dispensable for viability in *Caulobacter crescentus*, conditional upon the absence of Fur and the presence of a previously uncharacterized anionic sphingolipid, ceramide phosphoglycerate (**Fig. 7B**). LPS molecules form a robust permeability barrier based on 1) tight packing of the six saturated acyl chains of lipid A, and 2) a lateral network formed by the bridging of phosphate groups on lipid A or the core polysaccharide by divalent cations such as Mg^2+^ and Ca^2+^ (Nikaido, 2003). *Caulobacter* lipid A and core polysaccharide lack the phosphates that would participate in a lateral network with divalent cations (Smit et al., 2008). We propose that negative charges on CPG/CPG2 can provide this function in the *Caulobacter* OM, accounting for the observation that *cpgABC* and other sphingolipid synthesis genes are important for fitness even in non-stress conditions (**Fig. 5A** and Christen et al., 2011). Evidence that CPG/CPG2 contribute negative charge to the *Caulobacter* OM comes from studies of CAMP sensitivity. We previously observed that Spt is necessary for susceptibility to polymyxin B, but Sgt1 and Sgt2, which convert neutral ceramide to the anionic glycosphingolipid GSL-2, are not required (Stankeviciute et al., 2019).

This result was puzzling, because neutral ceramide was not expected to be a target for CAMP activity. Here we provide an explanation by showing that neutral ceramide is converted by CpgABC to a different anionic species, CPG2, and that this lipid, rather than LPS, is critical for colistin susceptibility.

### Inhibition of Fur-mediated gene expression is necessary to survive in the absence of lipid A

In contrast to Sphingomonads, the presence of cell surface sphingolipids is not sufficient for *Caulobacter* to survive in the absence of lipid A. Instead, the iron-responsive transcription factor Fur must also be deactivated. Both iron limitation (via growth in 2,2-dipyridyl) and excess available iron (due to a disruption in iron homeostasis in Δ deficient *Caulobacter*. These results imply that genes or processes regulated by Fur in complex with iron, rather than those regulated by iron independently of Fur, are the critical factors.

Fur controls iron homeostasis in *Caulobacter* by directly or indirectly regulating ∼120 genes in combination with iron (Leaden et al., 2018; da Silva Neto et al., 2009, 2013). A significant fraction of the Fur regulon, comprising 45 genes, is predicted to encode membrane proteins functioning in transport reactions or energy metabolism. *Caulobacter* Fur represses the transcription of genes predicted to mediate iron uptake, and it activates the expression of genes encoding iron-containing enzymes, including respiratory complexes containing Fe/S clusters or heme groups. Fur is linked to oxygen signaling in *Caulobacter* by activating the transcription of *fixK*, which mediates the response to hyxpoxia (Crosson et al., 2005). In addition, the Δ has a constitutively elevated level of intracellular oxidation, implicating Fur in the prevention of oxidative stress (Leaden et al., 2018).

Our *ctpA* suppressor screen retrieved mutations in *fur*, but not in genes whose transcription is activated by Fur. Thus, there is unlikely to be a singular Fur-regulated gene whose expression prevents the elimination of lipid A. Consistently, RB-Tnseq revealed that no transposon insertions in Fur-activated genes led to significantly increased fitness during challenge with CHIR-090. Since suppressor mutations would be more likely to cause loss than gain of function, we might not retrieve suppressors that work by increasing gene expression or activity. However, if there were a single Fur-repressed gene whose upregulation was critical for the ability to lose lipid A, then transposon insertions in this gene would be expected to reduce the fitness of CHIR-090-treated cells. Again, no individual gene fits this profile, but one caveat is that essential genes are excluded from RB-Tnseq analysis.

Based on our genetic data, mutations in *fur* could support the viability of lipid A-deficient *Caulobacter* via 1) downregulation of multiple Fur-activated genes, 2) upregulation of multiple Fur-repressed genes, and/or 3) activation of compensatory cellular stress responses. Since Fur regulates the expression of many OM and IM proteins, deletion of *fur* could alter envelope composition in a manner that renders lipid A non-essential. Alternatively, the transcriptional changes and oxidative stress which follow from *fur* deletion could activate a network of stress responses which together make it possible to eliminate lipid A.

### The search for principles governing lipid A essentiality

Hypotheses to explain the essential nature of lipid A include its chemical barrier function, the detrimental activation of stress responses when it is depleted, its role in OM protein biogenesis or function, and its mechanical role in resisting turgor pressure (Rojas et al., 2018; Zhang et al., 2013). *Caulobacter* is only the fourth LPS-bearing Gram-negative bacterium demonstrated to survive in the absence of lipid A, following *N. meningitidis*, *M. catarrhalis*, and *A. baumannii*. So far, however, no single theme has emerged to explain why this select and phylogenetically diverse group of Gram-negative species is capable of surviving without lipid A.

In *A. baumannii*, proteins which synthesize PG in lateral cell walls (the elongasome) are critical for the fitness of lipid A-deficient strains, suggesting that alterations in PG structure are needed to compensate for the OM’s loss of mechanical strength (Simpson et al., 2021). However, since elongasome components are essential for viability in wild-type *Caulobacter* (Christen et al., 2011), RB-Tnseq could not reveal their fitness effects in CHIR-090-treated cultures. Lipid A-deficient strains of *A. baumannii* consistently display increases in the expression of lipoproteins and the Lol pathway for lipoprotein transport to the OM (Boll et al., 2016; Henry et al., 2015). Two lipoprotein synthesis genes, *lgt* (*CCNA_00525*) and *lnt* (*CCNA_00050*), had markedly reduced fitness scores in CHIR-090-treated *Caulobacter* cultures compared to unstressed cultures, so OM lipoproteins may help to compensate for the absence of lipid A in diverse species.

*A. baumannii* Δ*lpxC* mutants have growth and morphological defects that are corrected when the growth rate is limited by environmental factors such as low temperature or nutrient limitation (Nagy et al., 2019), suggesting that one barrier to the elimination of lipid A is the rate of synthesis of alternative molecules to constitute the OM. Although Δ*fur* slows the growth of *Caulobacter* (**Fig. S1, Table S3**, and da Silva Neto et al., 2009), we found that slow growth in PYE medium at a reduced temperature was not sufficient to support the viability of *fur^+^ Caulobacter* depleted of LpxC (**Fig. S5A**), or of Δ*lpxC* Δ*fur* Δ*sspB* cells with *fur* expression restored (**Fig. S5B**). The Δ*fur* mutation therefore provides a benefit beyond decreasing the growth rate, which remains to be elucidated.

Our work suggests that possession of genes to produce anionic sphingolipids may provide certain bacteria with a unique pathway to the elimination of lipid A. In addition to facilitating OM remodeling, anionic sphingolipids can also underlie clinically important phenotypes, even in wild-type membranes that retain LPS, such as the susceptibility to CAMPs, which are used as a last line of defense against multidrug-resistant bacterial infections. Thus, functions traditionally attributed to lipid A may be performed wholly or in part by alternative lipids, underscoring the need to study lipid A functions in diverse species and to identify and functionally characterize novel lipids.

## Acknowledgments

The authors thank Charlie Huang, Morgan Price, Laura Herron and Henry Seaborn for experimental assistance. Cryoelectron microscopy data acquisition was performed using instrumentation of Lawrence Berkeley National Laboratory and the Bay Area CryoEM resource, maintained by Jonathan Remis and Daniel Toso respectively. Assistance in Linux computer maintenance was provided by Paul Tobias. D.R.G. thanks the International Centre for Cancer Vaccine Science project of the International Research Agendas program of the Foundation for Polish Science co-financed by the European Union under the European Regional Development Fund MAB/2017/03 at the University of Gdansk. Funding for this work was provided by National Science Foundation awards 1553004 and 2031948 to E.A.K. and 1615287 to K.R.R., and National Institutes of Health awards GM111066-01 and 1R01AI123820-01 to D.R.G. Cryoelectron tomography work was supported by the Laboratory Directed Research and Development Program of Lawrence Berkeley National Laboratory under U.S. Department of Energy Contract No. DE-AC02-05CH11231.

## Author contributions

J.J.Z., E.A.K., and K.R.R. conceived the project. Mass spectrometry experiments were performed and analyzed by S.H.Y., D.R.G., and Z.G. Electron cryotomography experiments were performed and analyzed by R.G. and K.M.D. RB-Tnseq experiments, sequencing, and data analysis were performed by J.J.Z and A.M.D., and K.R.R. All other experiments were performed and analyzed by J.J.Z., G.S, E.A.K., and K.R.R. The manuscript was written by J.J.Z., E.A.K, and K.R.R. with input from all authors.

## Methods

### Growth conditions

All *Caulobacter crescentus* strains were derived from NA1000 (Evinger and Agabian, 1977) and are listed in **Table S4**. Unless otherwise stated, *Caulobacter* was grown in peptone-yeast extract medium (PYE) (Ely, 1991) at 30°C. Solid media were prepared using Fisher agar (BP1423). PYE was supplemented with 0.3% xylose (PYEX) or 0.2% D-glucose (PYED) where indicated. When changing between inducing and non-inducing conditions, cells were washed twice with PYE medium lacking additional sugars before being released into or plated on medium supplemented with a different sugar. Counter-selection using *sacB* was performed using 3% sucrose. 100 µM 2,2’-dipyridyl was added to culture media to achieve low-iron conditions. Vanillic acid was added to PYE media at a final concentration of 0.5 mM to drive gene expression from the *vanA* promoter. Antibiotics added to PYE were used at the following concentrations (µg/mL) for liquid (L) or solid (S) medium: kanamycin, 5 (L), 25 (S); chloramphenicol, 1 (L/S); nalidixic acid, 20 (S); gentamycin, 25 (L), 5 (S); oxytetracycline, 1 (L), 2 (S); spectinomycin, 25 (L), 100 (S); hygromycin, 100 (L/S); streptonigrin 0.025 or 0.25 (L). *E. coli* was grown in lysogeny broth (10 g/L tryptone, 5 g/L yeast extract, 5 g/L NaCl) at 37°C, supplemented with antibiotics at the following concentrations (µg/mL) for liquid (L) or solid (S) medium: kanamycin, 30 (L), 50 (S); chloramphenicol, 20 (L), 30 (S); gentamycin, 15 (L), 20 (S); tetracycline, 12 (L/S); spectinomycin, 50 (L/S); hygromycin, 100 (L/S). Diaminopimelic acid (0.3 mM) was added to solid or media to support the growth of *E. coli* strain WM3064 (Dehio and Meyer, 1997).

### Plasmid construction

Plasmid descriptions are listed in **Table S5**. Primer sequences used for plasmid construction are listed in **Table S6**. *pZIK133*. The LpxC depletion vector was constructed by placing the *lpxC* coding region, C-terminally fused to a 3xFLAG tag (amino acid sequence: DYKDHDGDYKDHDIDYKDDDDK) followed by the *Caulobacter* ssrA tag (amino acid sequence: AANDNFAEEFAVAA), under control of the *xylX* promoter. The *xylX* promoter was amplified using the pJS14-PxylX and PxylX-lpxC R primers. The PxylX-lpxC F and lpxC-3xFLAG R primers were used to amplify *lpxC*. The C-terminal fusion was amplified from pAB6 using the lpxC-3xFLAG F and ssrA-pJS14 primers. The final plasmid was assembled via Gibson cloning into a BamHI/EcoRI-digested pJS14 backbone.

*pZIK134*. For the *lpxC* knockout construct, flanking homology regions were amplified using the primers lpxC UpF and lpxC UpR for the 5’-region, and lpxC DownF and lpxC DownR for the 3’-region. The 5’- arm included a 5’- SpeI site and a 3’- EcoRI site, and the 3’- arm included a 5’- EcoRI site and a 3’- SphI site. These fragments were digested with the indicated enzymes and ligated into SpeI/SphI-digested pNPTS138. This intermediate plasmid was linearized with EcoRI, and the EcoRI-digested *tetAR* cassette from pKOC3 was inserted to make the final construct.

*pZIK73* and *pZIK78*. For knockouts of *CCNA_01553* or *CCNA_00497*, flanking homology regions were amplified using the following primer pairs: pZIK73 5’- region (01553 UpF; 01553 UpR), pZIK73 3’- region (01553 DownF; 01553 DownR), pZIK78 5’- region (00497::hyg UpF; 00497::hyg UpR), pZIK78 3’- region (00497::hyg DownF; 00497::hyg DownR). For each construct, the 5’- arm included a 5’- SpeI site and a 3’- SmaI site, and the 3’- arm included a 5’- SmaI site and a 3’- EcoRI site. These fragments were digested with the indicated enzymes and ligated into SpeI/EcoRI-digested pNPTS138. The intermediate plasmids were linearized with SmaI, and the SmaI-digested *hyg* cassette from pHP45Ω constructs.

*pZIK80*, *pZIK81*, *pZIK82*, and *pZIK161*. For the knockouts of *CCNA_03733*, *CCNA_01068*, *CCNA_01055*, or *CCNA_00055*, flanking homology regions were amplified using the following primer pairs: pZIK80 5’- region (03733::hyg UpF; 03733::hyg UpR), pZIK80 3’- region (03733::hyg DownF; 03733::hyg DownR), pZIK81 5’- region (01068::hyg UpF; 01068::hyg UpR), pZIK81 3’- region (01068::hyg DownF; 01068::hyg DownR), pZIK82 5’- region (01055::hyg UpF; 01055::hyg UpR), pZIK82 3’- region (01055::hyg DownF; 01055::hyg DownR), pZIK161 5’- region (fur UpF; fur UpR), pZIK161 3’- region (fur DownF; fur DownR). Each 5’- arm included a 5’- SpeI site and a 3’- BamHI site, and each 3’- arm included a 5’- BamHI site and a 3’- EcoRI site. These fragments were digested with the indicated enzymes and ligated into SpeI/EcoRI-digested pNPTS138. The intermediate plasmids were linearized with BamHI, and the BamHI-digested hyg cassette from pHP45Ω-hyg was inserted to make the final constructs.

*pZIK172-174. CCNA_00497, CCNA_01553, or CCNA_03733* were placed under control of the *xylX* promoter on pXCERN-2, which integrates at the *xylX* promoter. The corresponding genes were initially cloned into pVCERN-2 before being moved into pXCERN-2. Genes were amplified with the following primer pairs: *CCNA_00497* (pVCERN-2 00497 F; pVCERN-2 00497 R), *CCNA_01553* (pVCERN-2 01553 F; pVCERN-2 01553 R), *CCNA_03733* (pVCERN-2 03733 F; pVCERN-2 03733 R). Primer sets replace the start codon with an NdeI site and add a SacI site after the stop codon. The corresponding gene fragment and pVCERN-2 were digested with NdeI and SacI and ligated together. An NdeI/MluI fragment was subsequently excised from each vector and moved to pXCERN-2 cut with the same enzymes.

*pZIK175*. *CCNA_00055* (*fur*) was placed under control of the *xylX* promoter on pXCERN-2, which integrates at the *xylX* promoter. *CCNA_00055* was initially cloned into pVCERN-2 before being moved into pXCERN-2. *CCNA_00055* was amplified using the Pvan-fur and fur-pVCERN primers, and this fragment was inserted into NdeI/SacI-digested pVCERN-2 via Gibson assembly. The NdeI/MluI fragment was subsequently excised and ligated into NdeI/MluI-digested pXCERN-2.

*pGS74 and pGS76.* For markerless deletions of *CCNA_01217* or *CCNA_01219*, 5□- and 3□-flanking homology regions, respectively, were amplified using the primer pairs EK1047/1048 and EK1049/1050 (*CCNA_01217*) and EK1055/1056 and EK1057/1058 (*CCNA_01219*). pNPTS138 was amplified with primers EK897/898, and vectors were constructed by Gibson assembly.

*pKR429*. For markerless deletion of *CCNA_01218*, 5□- and 3□- flanking homology regions, respectively, were amplified using the primer pairs 01218 up_fwd/01218 up_rev and 01218 down_fwd/01218 down_rev. pNPTS138 was digested with EcoRI and HindIII, and the vector was constructed by Gibson assembly.

*pEK406.* For complementing the deletion of *CCNA_01217* in LC-MS/MS experiments, *CCNA_01217-FLAG* was amplified using primers EK1357/1358. The PCR product was ligated into the NdeI/NheI sites of pVCHYC-5.

*pKR432*-*4*. For appending a C-terminal FLAG tag to the *CCNA_01218-20* open reading frames. The indicated genes were amplified from NA1000 genomic DNA using primer pairs Nde-01218/01218-Mlu, Nde-01219/01219-Mlu, Nde-01220/01220-Mlu. Fragments were digested using NdeI/MluI and ligated into pFLGC-1 digested with the same enzymes.

*pKR435.* For expressing *CCNA_01218-FLAG from the chromosomal vanA promoter. CCNA_01218-FLAG* was amplified from pKR432 using primers 01218-FLAG F/01218-FLAG R and inserted in NdeI-digested pVGFPC-2 by Gibson assembly.

*pKR436.* For expressing *CCNA_01219-FLAG from the chromosomal vanA promoter. CCNA_01219-FLAG* was amplified from pKR433 using primers 01219-FLAG F/01219-FLAG R and inserted in NdeI-digested pVGFPC-2 by Gibson assembly.

*pKR437.* For expressing *CCNA_01220-FLAG from the chromosomal vanA promoter. CCNA_01220-FLAG* was amplified from pKR434 using primers 01220-FLAG F/01220-FLAG R and inserted in NdeI-digested pVGFPC-2 by Gibson assembly.

*pKR438*. For expressing *CCNA_01217-FLAG from the chromosomal vanA promoter. CCNA_01217-FLAG* was amplified from pEK406 using primers 01217-FLAG F/01217-FLAG R and inserted in NdeI-digested pVGFPC-4 by Gibson assembly.

### Strain construction

Unless otherwise stated, plasmids were mobilized from *E. coli* into *C. crescentus* by conjugation. *E. coli* donors were counterselected by the addition of nalidixic acid, or when WM3064 was used as the donor, by omitting diaminopimelic acid from selection plates. Gene deletion or disruption was achieved by two-step homologous recombination using *sacB* counterselection (Ely, 1991).

#### CtpA depletion strain

In the CtpA depletion strain (KR3906, Shapland et al., 2011), regulated depletion of CtpA is achieved by expressing *ctpA::3xFLAG::ssrA* from a xylose-inducible promoter (Meisenzahl et al., 1997) on a high-copy plasmid in a Δ*ctpA* strain also lacking the proteolytic adaptor *sspB* (Levchenko et al., 2000). The native CtpA protein could not be depleted without an ssrA tag to target it for proteolysis. However, addition of this tag made CtpA proteolysis so rapid that xylose-dependent expression of CtpA-3xFLAG-ssrA did not support *Caulobacter* viability. Further deleting *sspB*, which encodes a proteolytic adaptor for ssrA-tagged substrates, reduced the basal rate of CtpA-3xFLAG-ssrA degradation enough to permit complementation in PYE medium containing xylose (PYEX).

#### LpxC depletion strain

The LpxC depletion strain KR4007 was constructed in a parallel manner to the CtpA depletion strain KR3906. pZIK133 was introduced to KR1499 (Δ*sspB*) by conjugation and selection on PYE/chloramphenicol. pZIK134 was conjugated into this intermediate strain, and colonies were selected on PYEX/chloramphenicol/oxytetracycline. After *sacB* counterselection on PYEX/sucrose/oxytetracycline, colonies were screened for chloramphenicol^R^ kanamycin^S^ on PYEX.

#### Stable ΔctpA or ΔlpxC strains

To generate stable Δ*ctpA* or Δ*lpxC* strains without covering plasmids, candidate suppressor genes identified by screening were disrupted in KR3906 or KR4007, respectively, using two-step homologous recombination while cultivating the cells on PYEX. Intermediate strains (sucrose^R^ hygromycin^R^ kanamycin^S^) were grown in liquid PYED without chloramphenicol to permit loss of the *ctpA* or *lpxC* covering plasmid, plated on PYED, and tested for chloramphenicol^S^. Absence of *ctpA* was confirmed using primers ctpA KO F and ctpA KO R, and absence of *lpxC* was confirmed using primers lpxC KO F and lpxC KO R. The genomes of Δ*ctpA* Δ*fur* Δ*sspB* (KR4102) and Δ*lpxC* Δ*fur* Δ*sspB* (KR4103) were resequenced and contained no additional mutations. Stable Δ*ctpA* or Δ*lpxC* strains were further modified by electroporation with purified plasmids (Gilchrist and Smit, 1991) to restore xylose-driven suppressor gene expression.

#### Unmarked deletions in genes for ceramide phosphoglycerate synthesi

Deletions in *CCNA_01217*, *CCNA_01218*, *CCNA_01219,* and *CCNA_01220* in NA1000 or KR4077 were made by conjugation of the appropriate pNPTS138-based plasmid, followed by selection on PYE/kanamycin/nalidixic acid. After overnight growth in PYE, cells were plated on PYEX/sucrose, and sucrose^R^ colonies were screened for kanamycin^S^. Colony PCR with the following primers was used to detect the deletion of the indicated chromosomal genes: *CCNA_01217,* EK S238/S239; *CCNA_01218*, EK S240/S241; *CCNA_01219*, EK S242/S243; *CCNA_01220*, EK S216/S217. Loci were sequenced with the indicated primers to ensure the accuracy of in-frame deletions. Unmarked deletions of *CCNA_01217*, *CCNA_01218*, *CCNA_01219,* and *CCNA_01220* were made in KR4091 by conjugation of KR4091 with WM3064 harboring the appropriate pNPTS138-based plasmids, followed by selection on PYEX/kanamycin medium omitting diaminopimelic acid. After growth overnight in PYEX, cells were plated on PYEX/sucrose, and sucrose^R^ colonies were screened for kanamycin^S^. Colony PCR with the following primers was used to detect the deletion of the indicated chromosomal genes: *CCNA_01217,* EK S238/S239; *CCNA_01218*, EK S240/S241; *CCNA_01219*, EK S242/S243; *CCNA_01220*, EK S216/S217. Loci were sequenced with the indicated primers to ensure the accuracy of in-frame deletions. Strains were screened for oxytetracycline^R^ and hygromycin^R^ to ensure that they maintained deletions of *lpxC* and *fur*, respectively.

#### Complementation of genes for ceramide phosphoglycerate synthesis

To complement deletions of *CCNA_01217-01220*, the following plasmids were introduced by conjugation to place the complementing gene under control of the chromosomal *vanA* promoter: Δ*CCNA_01217*, pEK406 (for LC-MS studies) or pKR438 (for growth and chemical sensitivity assays); Δ*CCNA_01218*, pKR435; Δ*CCNA_01219*, pKR436; or Δ*CCNA_01220*, pKR437. When introducing plasmids into strains capable of *lpxC* depletion (based on KR4091), plasmids were delivered from WM3064 to avoid the use of multiple antibiotics for selection/counterselection. Correct integration of plasmids at the *vanA* locus was confirmed by colony PCR using primers RecUni-1 and RecVan-2 (Thanbichler et al., 2007).

#### Suppressor selection

KR3906 was grown to full density in PYEX. 300 µL of culture was transferred onto an open, sterile Petri dish and mutagenized in a UV Stratalinker 1800 (Stratagene) with 30,000 µJ of energy. Mutagenized cells were plated on PYED. Recovered colonies were passaged in liquid PYED overnight to allow loss of the covering plasmid, and samples were streaked onto PYED. Isolated colonies were screened for chloramphenicol sensitivity. Chlor^S^ isolates were grown in PYE and saved at −80°C in 10% dimethylsulfoxide. Loss of *ctpA* was confirmed via PCR using the primers ctpA KO F and ctpA KO R, which anneal to the interior of the open reading frame.

#### Genome resequencing

Strains were grown to full density in PYE, and genomic DNA was extracted using the Quick-DNA Miniprep Kit (Genesee) or the DNeasy Blood & Tissue Kit (Qiagen). Genomic DNA was submitted to the UC Berkeley Functional Genomics Laboratory, where libraries were prepared using a PCR-free protocol with multiplexing (http://qb3.berkeley.edu/gsl/). Samples were sequenced at the UC Berkeley Vincent J. Coates Genomics Sequencing Laboratory using a 300PE or 150PE MiSeq v3 run. Genomic sequencing data were analyzed for variants using the Galaxy platform at usegalaxy.org (Afgan et al., 2016). Adapter sequences were removed using Cutadapt, and sequences were aligned to the NA1000 genome (Marks et al., 2010) using Bowtie2. FreeBayes was used to analyze the BAM files for variants. Variants with quality scores below 300 were discarded as noise.

#### Growth and viability assays

For plate assays, strains were grown to OD_660_ = 0.2-0.5 in permissive media, washed twice in PYE medium with no additions, and diluted to OD_660_ = 0.1. 10 μl drops of ten-fold serial dilutions were pipetted onto permissive and nonpermissive media. Plates were incubated for 3 days at 30°C, and images are representative of at least three independent trials. For end-point growth assays in liquid media, strains were grown in permissive media to OD_660_ = 0.2-0.5. After washing in PYE medium without additions, cells were resuspended at OD_660_ = 0.01 in permissive and nonpermissive media. OD_660_ values were measured after 24 h growth at 30°C.

#### Disc diffusion assays of chemical sensitivity

Cultures were grown to mid-exponential phase (OD_660_ 0.2-0.5), and an amount of cells equivalent to 250 µl of culture at OD_660_ = 0.2 was added to 4 ml of PYE swarm agar (0.3% w/v agar) pre-warmed to 42°C. Swarm agar containing bacteria was spread onto solid PYE and allowed to set. Antibiotics or detergents (10 μl each) were added to sterile Whatman filter disks and allowed to dry in a fume hood before discs were placed onto swarm agar surfaces. Plates were incubated upright at 30°C for 24 hours. The diameters of the zones of clearing or haze were measured, and the diameter of the disk (6 mm) was subtracted from all measurements to yield the reported values. The total amount of antibiotic or detergent added to each disk is as follows: kanamycin (100 µg), rifampicin (100 µg), vancomycin (1 mg), CHIR-090 (100 µg, APExBIO), bacitracin (50 µg), TWEEN 20 (10 µL of 10% solution), Triton X-100 (10 µL of 10% solution), sodium dodecyl sulfate (10 µL of 10% solution). Tests using CHIR-090 used one quarter of the standard amount of cells to reduce growth haze. For strains overexpressing genes integrated at the *vanA* locus, uninduced cells were grown in PYE/kanamycin or PYE/gentamicin, and aliquots were plated in PYE swarm agar on PYE medium. Induced cells were grown in PYE/kanamycin or PYE/gentamicin containing 0.5 mM vanillate before plating in/on PYE medium containing 0.5 mM vanillate. 100 µM 2,2’-dipyridyl was included in media for testing chemical sensitivity in iron-restricted conditions.

#### Streptonigrin sensitivity

Isolated colonies of the indicated strains were grown in PYE medium to OD_660_ = 0.2-0.5 and diluted to OD_660_ = 0.01. The diluted culture was aliquoted into separate tubes, which received 0.025 μg/ml, 0.25 μg/ml, or no streptonigrin (SNG). After 24 h of growth at 30°C, OD_660_ values were measured, and optical density ratios (0.25 μg/ml SNG/no addition and 0.025 μg/ml SNG/no addition) were calculated as a measure of growth inhibition.

#### Limulus amebocyte lysate (LAL) assay

The ToxinSensor Chromogenic LAL Endotoxin Assay kit (GenScript) was used to determine endotoxin units/mL of culture. Cells were grown to mid-exponential phase (OD_660_ 0.2-0.5), washed twice with non-pyrogenic LAL reagent water, and normalized in this water to OD_660_ = 0.1. Cell suspensions were serially diluted in non-pyrogenic water and analyzed according to manufacturer’s instructions.

#### Extraction and visualization of LPS and lipid A species

For visualizing LPS species from whole-cell lysates, cells were harvested after overnight growth in the indicated medium. All cultures were normalized by OD_660_, pelleted, and resuspended to 100 µL in 1x tricine loading buffer (100 mM Tris-HCl pH 6.8, 1% sodium dodecyl sulfate (SDS), 20% glycerol, 0.02% Coomassie G-250, 1% 2-mercaptoethanol). Proteinase K (125 ng/µL) was added, and samples were incubated overnight at 55°C. Lysates were boiled 5 min, and equal volumes (10% of each sample) were analyzed by gel electrophoresis.

Hot aqueous-phenol LPS extractions were adapted from Westpahl and Jann (Davis and Goldberg, 2012; Westphal and Jann, 1965). 1 mL of culture at OD_660_ = 0.75 was pelleted and resuspended in 200 µL 1x tricine loading buffer. Suspensions were boiled for 15 min and cooled to room temperature. 5 µL of 20 mg/mL Proteinase K (Thermo) was added to each sample before incubation at 55°C for three hours. Suspensions were mixed with 200 µL ice-cold Tris-saturated phenol, vortexed, and incubated at 65°C for 15 minutes before being cooled to room temperature. 1 mL diethyl ether was added to each sample before vortexing and spinning for 10 minutes in a table-top centrifuge at 16,000 x *g*. The bottom blue layer was removed to a fresh tube, and the extraction was repeated on the blue layer starting from the phenol step. 200 µL 2x tricine loading buffer was added to each sample before gel electrophoresis.

Rough LPS was extracted by the method of Darveau and Hancock (Darveau and Hancock, 1983), modified as described (Hershey et al., 2019), beginning with 50 ml PYE cultures grown to OD_660_ = 0.85. Cultures were centrifuged, and cell pellets were resuspended in 2 ml 10 mM Tris-HCl (pH 8.0) containing 2 mM MgCl_2_. Samples were sonicated (Qsonica Q500) on ice for 5 min at 20% amplitude, in cycles of 10 sec on/20 sec off so that fewer than 5% of cells remained intact. DNase I and RNase A were added to final concentrations of 100 μg/ml and 25 μg/ml, respectively, and lysates were incubated at 37°C for 1 hour. Additional DNase I and RNase A were added to reach final concentrations of 200 μg/ml and 50 μg/ml, respectively, and lysates were incubated for 1 hour at 37°C. SDS and EDTA were added to achieve final concentrations of 2% and 100 mM, respectively, and lysates were incubated for 2 h at 37°C before centrifugation (30 min at 50,000 x *g*, 30 min, 4°C). Proteinase K (50 μg/ml) was added to each supernatant, followed by incubation for 2 h at 60°C. LPS was precipitated by the addition of 2 volumes of ice-cold 0.375 M MgCl_2_/95% ethanol and collected by centrifugation (12,000 x *g*, 15 min, 4°C). Precipitates were resuspended in 3.3 ml 10 mM Tris-HCl (pH8.0)/2% SDS/100 mM EDTA and incubated with shaking overnight at 37°C. Rough LPS was reprecipitated using 2 volumes ice-cold 0.375 M MgCl_2_/95% ethanol and collected by centrifugation (12,000 x *g*, 15 min, 4°C). Precipitates were suspended in 10 mM Tris-HCl (pH 8.0) and centrifuged (200,000 x *g*, 2 h, 4°C). After removal of the supernatant by pipetting, LPS pellets were resuspended in 1 mll 1x tricine loading buffer, and 5 μl were analyzed by gel electrophoresis.

Free lipid A was extracted by the Caroff method (El Hamidi et al., 2005), modified as described (Leung et al., 2017), starting with 10 ml of PYE culture grown to OD_600_ = 0.6. Cultures were divided into multiple tubes and centrifuged at 14,000 x *g* for 2 min. In a fume hood, cell pellets from each culture were resuspended, combined, and transferred to a gasketed microcentrifuge tube using 250 μl 70% (v/v) isobutyric acid + 150 μl 1 M ammonium hydroxide. Samples were incubated in a boiling water bath in a fume hood for 1 h, with vortexing every 15 min. Samples were cooled on ice and centrifuged at 2000 x *g* for 15 min. In a fume hood, supernatants (∼400 μl) were transferred to new gasketed tubes, each containing 400 μl endotoxin-free water. Small holes were punched in the gasketed caps using a syringe needle before the samples were frozen in liquid nitrogen and lyophilized overnight. Methanol (1 ml) was added, and samples were sonicated in a water bath for 5 min. Samples were centrifuged at 10,000 x *g* for 5 min, and methanol was aspirated. The methanol wash was repeated before lipids were solubilized in 190 μl 3:1.5:0.25 v/v/v chloroform:methanol:endotoxin-free water. After vortexing, samples were centrifuged at 8,000 x *g* for 5 min. Supernatants were transferred to fresh gasketed tubes, and extracts were dried under a stream of nitrogen before analysis by mass spectrometry (see below) or gel electrophoresis. Samples for gel electrophoresis were resuspended using 100 μl 1x tricine loading buffer, and 10 μl of each sample was analyzed.

All lipid samples were analyzed on 16.5% Mini-PROTEAN Tris-Tricine gels (Bio-Rad). Carbohydrates were stained using Pro-Q Emerald 300 Lipopolysaccharide Gel Stain Kit (Molecular Probes; P20495) per manufacturer’s instructions. For Western blot analysis of S-LPS, equal numbers of cells grown in PYE with appropriate additions were pelleted, resuspended in 1x SDS loading buffer, and boiled before analysis on 12% polyacrylamide gels and transfer to Immobilon-P PVDF membranes. Blots were probed with α-S-LPS (1:20,000) (Walker et al., 1994) and horseradish peroxidase-conjugated anti-rabbit antibodies (1:5000) and analyzed using Western Lightning. Stained lipid species were visualized using a Bio-Rad Gel Doc XR.

#### High performance liquid chromatography-tandem mass spectrometry (HPLC-MSMS) of lipid A extracts

All samples for HPLC-electrospray ionization tandem mass spectrometry were generated by the modified Caroff extraction protocol described above. Each extract was initially dissolved in 100 µL 1:2 chloroform: methanol before dilution 1:10 with methanol for analysis. A 2-5 µL aliquot of each solution was injected onto a Phenomenex Jupiter C4 column (2 x 50 mm, 5 µm, 300 Å) for HPLC-MSMS analysis with a Waters Acquity UPLC system coupled to a Thermo LTQ-Orbitrap Velos Pro mass spectrometer, which was equipped with an atmospheric pressure electrospray ionization source. For lipid detection, the HPLC-MSMS analyses were carried out with full-mass detection over a mass range of *m/z* 250 to 2000 in the Fourier transform MS mode, with negative-ion detection. The mass resolution was 60,000 FWHM @ *m/z* 400. Fragmentation product ion masses of the three most intense precursor ions were measured in the ion trap or orbitrap (7500 resolution) mass analyzer using stepped collision-induced dissociation (35% of the normalized collision energy) or Higher energy collision-induced dissociation (35% of the normalized collision energy) activation energies. During data acquisitions, real-time mass calibration was applied with *m/z* 283.26454 as the lock mass for negative-ion detection. The mobile phase for separation was (A) 1 mM ammonium acetate solution and (B) 90% (1:1 acetonitrile/propanol)/10% water/1 mM ammonium acetate as the binary solvents for the 16-minute gradient elution: 0 to 10 min, 30% to 100%B; 10 to 12 min, 100% B and 12 to 12.1 min at 30% B, followed by column equilibration at 30% B from 12.1 to 16 min. The column flow rate was 0.35 mL/min and the column temperature was maintained at 40°C.

#### Lipid A structure analysis

MALDI-TOF MS was used to screen lipid extracts. To check structures, tandem MS and ancillary separation techniques were required. These are described below. HPLC-MSMS (above) describes the generation of data for structure determinations in Fig. S3. Notably, the triple deletion strain Δ*ctpA* Δ*fur* Δ*sspB* contained no lipid A with sugars at the terminal (1 and 4′) positions but rather contained phosphates, as found in canonical lipid A structures. The Δ*lpxC* Δ*fur* Δ*sspB* strain contained an ion at 1412 *m/z*, the structure of which remains unclear. The HPLC-MSMS data of this ion showed no loss of phosphate, as seen in Δ*ctpA* Δ*fur* Δ*sspB*, nor loss of sugars, as seen for the NA1000, Δ*sspB*, or Δ*fur* Δ*sspB* strains. The fragmentation pattern strongly suggested that something other than lipid A was responsible for the ion at 1412 *m/z*. Given that cardiolipin is a common microbial membrane lipid, we carried out HILIC-MS (described below) with cardiolipin and lipid A standards. Both standards were retained by HILIC, as expected for hydrophobic molecules, but extracts from the Δ*lpxC* Δ*fur* Δ*sspB* mutant showed no ions at all, suggesting that the species at 1412 *m/z* is not hydrophobic enough to be retained. Regrettably, there remains no structure identified for the ion at 1412 *m/z*. Generally, structure analysis was conducted manually according to our prior effort in this field (Yoon et al., 2016).

#### Hydrophobic interaction liquid chromatography-mass spectrometry (HILIC-MS)

A 10-µL aliquot of each solution was injected into a Waters Atlantis HILIC column (4.6 mm x 150 mm, 5 µm) to run LC-MS on a Water Acquity UPLC system coupled to a Thermo LTQ-Orbitrap Velos Pro mass spectrometer, which was equipped with an atmospheric pressure electrospray ionization source. For lipid detection, the HILIC-MS runs were carried out with full-mass detection over a mass range of *m/z* 80 to 2000 in the Fourier transform MS mode, with positive-ion and negative-ion detection, respectively, in two rounds of LC injections. The mass resolution was 60,000 FWHM @ *m/z* 400. During data acquisitions, real-time mass calibration was applied with *m/z* 391.28426 as the lock mass for positive-ion detection and with *m/z* 112.98563 as the lock mass for negative-ion detection. The mobile phase of HILIC was (A) 20 mM ammonium acetate solution (pH adjusted to 4.0 with acetic acid) and (B) methanol as the binary solvents for gradient elution: 0-4 min, 99% B; 4 to 12.5 min, 99% to 20% B and 12.5 to 15 min at 20% B, followed by column equilibration at 99% B for 5 min between injections. The column flow rate was 0.4 mL/min and the column temperature was maintained at 40°C.

#### Differential interference contrast microscopy

Cells were immobilized on agarose pads (1% w/v in reverse osmosis-purified water). Images were taken using a Zeiss EC Plan-Neofluar 100x/1.3 Oil M27 objective on a Zeiss AxioImager M1 microscope with a Hamamatsu Digital CCD Camera (C8484-03G01). Images were acquired using iVision software and processed using ImageJ.

#### CryoEM imaging and tomographic processing

Cultures (5 mL) of KR4000, KR4102, KR4103, and KR3906 grown to OD_660_ 0.2-0.5 were centrifuged (4°C, 16,000 x *g*, 15 minutes), and cell pellets were resuspended in 50 µL PYE. For KR3906, cells grown in PYEX were washed twice with PYE, released into PYED at OD_660_ = 0.02, and incubated for 12 hours before harvest. 3 µL of cell suspension, mixed 1:1 with Fiducial markers (10-nm gold particles conjugated to Protein A; Aurion) was applied to glow-discharged quantifoil grids (R2/2) and frozen in liquid ethane using an automatic plunge freezing device (Vitrobot, FEI. 12°C, 8-12s blot time, blot force 8, humidity 100%).

Grids of KR4000 and KR4103 were imaged on a Jeol3100 cryoTEM operating at 300kV with in column omega energy filter and K2 direct electron camera. Grids of KR4102 and KR3906 were imaged on a Krios Cryo TEM (FEI) operating at 300kV with post column energy filter (Quantum, GATAN) and K2 direct electron camera. All data were collected with the automatic data collection program serialEM (Mastonarde, 2005). Square overview images were acquired using a defocus of 80-100 microns at a nominal magnification of 3600-6500x (Krios) or 1200x (Jeol) using the polygon montage operation (specimen pixel size: 33-67Å). Beam intensity was set to 8e^-^/px/s over an empty hole and exposure times ranged from 2-5s depending on ice thickness. Bidirectional tomographic tilt series were collected from ±60° using a defocus of 6-8 µm and at a magnification which provided specimen pixel size of 4-7 Å. Total dose of the tilt series were kept between 60-90 e^-^/Å^2^. All tilt series images were collected in movie mode and the frames aligned using MotionCor2 (Zheng et al., 2017). Aligned frames were compiled into stacks and processed using IMOD (Kremer et al., 1996). Contrast of resulting tomograms was enhanced using a non-linear anisotropic diffusion filter (Frangakis and Hegerl, 2001) and manually segmented using the 3D visualization program AMIRA (ThermoFisher).

#### RB-Tnseq analysis

A 1 ml aliquot of the RB-Tnseq library in NA1000 (Price et al., 2018) was thawed and grown to OD_660_ = 0.65 in 25 ml PYE medium with kanamycin. Aliquots of this culture were saved for sequencing of pre-challenge barcodes, or were diluted to OD_660_ = 0.02 in PYE medium (set8IT011, set8IT023, and set8IT035) or PYE medium with 2 μ (set8IT012, set8IT024, and set8IT036). Cultures were grown for 9 hours at 30°C as described (Price et al., 2018) before cells were harvested and post-challenge barcodes were sequenced. Gene fitness (f) and significance (t) scores were calculated as described (Wetmore et al., 2015). Candidate genes examined in this study (*CCNA_01217-01220*) had fitness scores between −1.5 and −3.7, with signficance scores between −3.0 and −8.4, for individual trials of library growth in PYE + CHIR-090.

#### Extraction and liquid chromatography-tandem mass spectrometry (LC-MS/MS) of sphingolipids

*Caulobacter* strains were grown overnight with or without 0.5 mM vanillate (5 ml), and lipids were extracted by the method of Bligh and Dyer (Bligh and Dyer, 1959). Cells were harvested and resuspended in 1 ml of water, 3.75 ml of 1:2 (v/v) chloroform: methanol was added, and the samples were mixed by vortexing. Chloroform (1.25 ml) and water (1.25 ml) were added sequentially with vortexing to create a two-phase system and the samples were centrifuged at 200 x *g* for 5 minutes at room temperature. The bottom, organic phase was transferred to a clean glass tube with a Pasteur pipette and washed twice in “authentic” upper phase. Subsequently, the organic phase containing lipids was collected and dried under argon. Our methods for lipid analysis by normal phase LC/ESI–MS/MS have been described (Guan et al., 2014). Briefly, normal phase LC was performed on an Agilent 1200 Quaternary LC system equipped with an Ascentis Silica HPLC column, 5 µm, 25 cm × 2.1 mm (Sigma-Aldrich, St. Louis, MO) as described. The LC eluent (with a total flow rate of 300 µl/min) was introduced into the ESI source of a high resolution TripleTOF5600 mass spectrometer (Applied Biosystems, Foster City, CA). Instrumental settings for negative ion ESI and MS/MS analysis of lipid species were as follows: ion spray voltage (IS) = −4500 V; curtain gas (CUR) = 20 psi; ion source gas 1 (GSI) = 20 psi; declustering potential (DP) = −55 V; and focusing potential (FP) = −150 V. The MS/MS analysis used nitrogen as the collision gas. Data analysis was performed using Analyst TF1.5 software (Applied Biosystems, Foster City, CA).

### Data Availability Statement

Sequence data that support the findings of this study are openly available in the Sequence Read Archive at https://www.ncbi.nlm.nih.gov/sra, under BioProject ID PRJNA526705, with specific NCBI BioSample accession numbers listed in **Table S3**. RB-Tnseq data are accessible at https://fit.genomics.lbl.gov/, with set and index numbers listed under RB-Tnseq analysis.

## Supplemental Information Titles and Legends

**Figure S1, related to Fig. 1:** Morphology, growth, and complementation of strains in which suppressor mutations permit the loss of *ctpA.* (A) Isolates from the Δ*ctpA* suppressor screen show variations in morphology. DIC images of selected suppressor isolates confirmed to have lost the *ctpA* covering plasmid. Putative suppressor mutations identified by whole-genome resequencing are indicated. Scale bar, 3 μm. (B) Growth curves of the indicated strains in PYE showing (i or iii) OD_660_ and (ii or iv) colony-forming units (CFU) per mL (mean ± S.D., *N*=3). (C) Viability assays of Δ*ctpA* suppressor mutants, each harboring a vector for xylose-driven expression of the corresponding suppressor gene or the *cerulean* gene as a control. Kanamycin was included in media to retain expression vectors.

**Figure S2, related to Fig.1:** A subset of *ctpA* suppressor mutations impair or block S-LPS production. (A) α-S-LPS-probed immunoblot and (B) Pro-Q Emerald 300-stained gel of Proteinase K-treated whole-cell lysates of the indicated strains. (C) Complementation of O-antigen biosynthesis using plasmid-borne genes driven by a xylose-inducible promoter. Pro-Q Emerald 300-stained polyacrylamide gel of Proteinase K-treated whole-cell lysates of the indicated strains grown in either PYED (D) or PYEX (X). Samples were normalized by OD_660_. *** = S-LPS, ** = putative full-length lipid A-core polysaccharide, * = putative incomplete lipid A-core species in cells lacking *manC* activity (*CCNA_03733*).

**Figure S3, related to Figure 2:** Δ*ctpA* and Δ*lpxC* strains with suppressor mutations lack wild-type lipid A. Tandem mass spectrometry (MSMS)-derived structures of lipid A from the indicated strains. Lipid extraction and MSMS analysis were performed using the same protocols for all strains.

**Movie S1, related to Figure 2.** Tomogram of *Caulobacter crescentus* NA1000 grown to mid-exponential phase in PYE.

**Movie S2, related to Figure 2.** Tomogram of Δ*ctpA* Δ*sspB* Δ*fur* grown to mid-exponential phase in PYE.

**Movie S3, related to Figure 2.** Tomogram of Δ*lpxC* Δ*sspB* Δ*fur* grown to mid-exponential phase in PYE.

**Movie S4, related to Figure 2.** Tomogram of Δ*ctpA* Δ*sspB* + pJS14-P*_xylX_*-*ctpA-3xFLAG-ssrA* grown to mid-exponential phase in PYEX, washed twice with PYE, released into PYED at OD_660_ 0.02, and incubated for 10 hours before harvest.

**Figure S4, related to Figure 6:** The total ion chromatogram of the *C. crescentus* lipidome shows the major lipid species present. DAG: diacylglycerol; MHDAG: mono-hexosyl diacylglycerol; FA: fatty acids; PG: phosphatidylglycerol; HexA-DAG: hexuronic acid-diacylglycerol; CPG: ceramide phosphoglycerate.

**Figure S5, related to Figure 3.** Slow growth at a reduced temperature in rich medium is insufficient for viability of Δ*lpxC* strains. (*A*) Viability of the LpxC depletion strain grown in inducing (PYEX) or depleting (PYED) conditions, at the indicated temperatures. (*B*) Viability of the stable Δ*lpxC* Δ*fur* Δ*sspB* strain harboring a P*xyl*-*fur* plasmid, grown in noninducing (PYED) or inducing (PYED) conditions, at the indicated temperatures. Cells were plated from 10-fold serially diluted suspensions normalized to OD_660_ = 0.1. Plates were incubated for 3 days (30°C) or 6 days (22°C) and are representative of at least three independent trials. Plates in *B* included kanamycin to retain the *fur* expression vector.

## Supplementary Tables

**Table S1, related to Figure 1:** Single-nucleotide polymorphisms and indels in _Δ_ctpA suppressors^a^.

**suppressors<sup>a</sup>**
| Strain | Sequence position<sup>b</sup> | Gene<sup>c</sup> | Annotation | Base Change | Amino Acid Substitution |
| --- | --- | --- | --- | --- | --- |
| S3 | 1668695 | 01553 | Undecaprenyl-phosphate beta-N-acetyl-D-fucosaminephosphotransferase | G>T | K54N |
|  | 2383967 | 02235 | SGNH hydrolase family protein | G>A | G109D |
| S8 | 3492550 | 03316 | UDP-N-acetylglucosamine 4,6-dehydratase/UDP-D-quinovosamine 4-dehydrogenase | AG>ATG | E487Frameshift |
| S16 | 58496 | 00055 | Ferric uptake regulation protein | A>T | L38Q |
| S21 | 515085 | 00497 | Putative rhamnosyl transferase | G>A | G263D |
|  | 1378399 | 01250 | FecCD-family transporter protein | GCC>GC | A340Frameshift |
| S32 | 1157376 | 01056 | Methyltransferase | A>AG | S31Frameshift |
|  | 2164248 | 02016 | nuoMNADH-quinone oxidoreductase chain M | C>T | M7I |
|  | 2949436 | 02792 | TonB-dependent outer membrane receptor | A>G | S516G |
| S36 | 3901011 | 03733 | Mannose-1-phosphate guanylyltransferase | C>T | W279Stop |
| S38 | 726779 | 00669 | Glycosyltransferase family 99 protein WbsX | CTG>CG | Q476Frameshift |
| S40 | 378822 | 00362 | Zinc uptake regulation protein | A>T | C156S |
|  | 2868272 | - | - | G>C | - |
|  | 3913883 | 03744 | dTDP-glucose 4,6-dehydratase | T>A | I269F |
| S43 | 1377138 | 01249 | ABC-transporter substrate binding protein | C>T | A169V |
|  | 1668861 | 01553 | Undecaprenyl-phosphate beta-N-acetyl-D-fucosaminephosphotransferase | CC>GA | P110E |
|  | 2976030 | 02820 | TadG-family protein | C>T | Silent |
| S44 | 727833 | 00669 | Glycosyltransferase family 99 protein WbsX | A>C | L125R |
| S47 | 537654 | 00524 | Conserved hypothetical cytosolic protein | A>AG | L369Frameshift |
|  | 1173251 | 01068 | Glycosyltransferase | GCC>GC | R293Frameshift |
| S52 | 514656 | 00497 | Putative rhamnosyl transferase | T>G | L120R |
|  | 810384 | 00752 | 3-hydroxybutyryl-CoA dehydrogenase | T>A | Stop>Y |
|  | 3797776 | - | - | T>C | - |
| S53 | 58379 | 00055 | Ferric uptake regulation protein | G>T | S77Stop |
|  | 1778110 | 01656 | Endonuclease/exonuclease/phosphatase family protein | A>C | L4R |
|  | 3900664 | 03733 | Mannose-1-phosphate guanylyltransferase | G>A | Q395Stop |
| S54 | 1377127 | 01249 | ABC-transporter substrate binding protein | CGG>CG | G166Frameshift |
|  | 3492771 | 03316 | UDP-N-acetylglucosamine 4,6-dehydratase/UDP-D-quinovosamine 4-dehydrogenase | G>A | E561K |
| S57 | 1492703 | 01378 | Protein-L-isoaspartate O-methyltransferase | G>GC | G59Frameshift |
|  | 2487074 | 02347 | Phosphomannomutase/phosphoglucomutase | G>A | G266D |
| S111 | 295119 | 00283 | 2,3,4,5-tetrahydropyridine-2,6-dicarboxylate N-succinyltransferase | A>T | L284Q |
|  | 1173965 | 01068 | Glycosyltransferase | TGC>T | R55Frameshift |
| S112 | 311705 | 00297 | Two-component response regulator | C>T | W64Stop |
|  | 315570 | 00301 | Phosphotransferase family protein | T>G | I319S |
|  | 2487307 | 02347 | Phosphomannomutase/phosphoglucomutase | G>A | E344K |
|  | 3569520 | 03399 | Flavin prenyltransferase UbiX | C>T | A159T |

<sup>a</sup>In addition to the indicated mutations, each strain is also $\Delta ctpA \Delta sspB$ .
<sup>b</sup>Sequence positions refer to the genome of *Caulobacter crescentus* NA1000 (NC\_011916.1).
<sup>c</sup>The prefix for each gene number is CCNA\_.

**Table S2, related to Figure 3:** Single-nucleotide polymorphisms and indels identified in ΔlpxC suppressors^a^.

| Strain | Sequence position<sup>b</sup> | Gene<sup>c</sup> | Annotation | Base Change | Amino Acid Substitution |
| --- | --- | --- | --- | --- | --- |
| S1 | 58446 | 00055 | Ferric uptake regulation protein Fur | A>T | Y55N |
|  | 4000704 | 03835 | 3-oxoacyl-(Acyl-carrier-protein) synthase | C>T | G396S |
| S5 | 58436 | 00055 | Ferric uptake regulation protein Fur | A>T | V58E |

<sup>a</sup>In addition to the indicated mutations, each strain is also $\Delta lpxC \Delta sspB \Delta 00497$ .
<sup>b</sup>Sequence positions refer to the genome of *Caulobacter crescentus* NA1000 (NC\_011916.1).
<sup>c</sup>The prefix for each gene number is *CCNA\_*.

**Table S3, related to Figure 3:** Growth rates in PYE medium at 22°C.

|  | Doubling time +/- standard deviation (hours)<sup>a</sup> |  |
| --- | --- | --- |
| Strain | PYE 30°C | PYE 22°C |
| NA1000 | 1.61 +/- 0.04 | 2.50 +/- 0.17 |
| $\Delta sspB$ | 1.51 +/- 0.04 | 2.44 +/- 0.09 |
| $\Delta fur \Delta sspB$ | 1.77 +/- 0.08 | 3.16 +/- 0.21 |

<sup>a</sup>Doubling times were calculated from optical density measurements (660 nm) during exponential growth.

**Table S4, related to STAR Methods:** Strains used in this study.

| Strain Number | Description | Note | NCBI BioSample Accessions | Reference |
| --- | --- | --- | --- | --- |
| KR4000 | Wild-type *Caulobacter* NA1000 |  |  | (Evinger and Agabian, 1977) |
| KR3180 | NA1000 pJS14 |  |  | This study |
| KR1499 | $\Delta sspB::aadA$ | | | (Shapland et al., 2011) |
| KR3877 | $\Delta CCNA\_00497::hyg$ | | | This study |
| KR4198 | $\Delta CCNA\_00497::hyg$ pJS14 | | | This study |
| KR3871 | $\Delta CCNA\_01553::hyg$ | | | This study |
| KR4197 | $\Delta CCNA\_01553::hyg$ pJS14 | | | This study |
| KR4076 | $\Delta fur::hyg$ | | | This study |
| KR4199 | $\Delta fur::hyg$ pJS14 | | | This study |
| KR3953 | $\Delta CCNA\_00497::hyg \Delta sspB::aadA$ | | | This study |
| KR4115 | $\Delta CCNA\_01055::hyg \Delta sspB::aadA$ | | | This study |
| KR4116 | $\Delta CCNA\_01068::hyg \Delta sspB::aadA$ | | | This study |
| KR3954 | $\Delta CCNA\_01553::hyg \Delta sspB::aadA$ | | | This study |
| KR3955 | $\Delta CCNA\_03733::hyg \Delta sspB::aadA$ | | | This study |
| KR4077 | $\Delta fur::hyg \Delta sspB::aadA$ | | | This study |
| KR4153 | $\Delta sspB::aadA$ pXCERN-2 | | | This study |
| KR4154 | $\Delta CCNA\_00497::hyg \Delta sspB::aadA$ pXCERN-2 | | | This study |
| KR4155 | $\Delta CCNA\_01553::hyg \Delta sspB::aadA$ pXCERN-2 | | | This study |
| KR4156 | $\Delta CCNA\_03733::hyg \Delta sspB::aadA$ pXCERN-2 | | | This study |
| KR4157 | $\Delta fur::hyg \Delta sspB::aadA$ pXCERN-2 | | | This study |
| KR4158 | $\Delta CCNA\_00497::hyg \Delta sspB::aadA$ pZIK172 | | | This study |
| KR4159 | $\Delta CCNA\_01553::hyg \Delta sspB::aadA$ pZIK173 | | | This study |
| KR4160 | $\Delta CCNA\_03733::hyg \Delta sspB::aadA$<br>pZIK174 | | | This study |
| KR4161 | $\Delta fur::hyg \Delta sspB::aadA$ pZIK175 | | | This study |
| KR4269 | $\Delta fur::hyg \Delta sspB::aadA$ pZIK197 | | | This study |
| KR3906 | $\Delta ctpA::tetAR \Delta sspB::aadA$ pAB6 | Restock of KR2423<br>from Shapland 2011 | | (Shapland et al., 2011) |
| KR4111 | $\Delta ctpA::tetAR \Delta sspB::aadA$<br>$\Delta CCNA\_00497::hyg$ pAB6 | | SAMN12568762 | This study |
| KR4112 | $\Delta ctpA::tetAR \Delta sspB::aadA$<br>$\Delta CCNA\_01553::hyg$ pAB6 | | SAMN12568763 | This study |
| KR4092 | $\Delta ctpA::tetAR \Delta sspB::aadA$<br>$\Delta CCNA\_03733::hyg$ pAB6 | | SAMN12568761 | This study |
| KR4090 | $\Delta ctpA::tetAR \Delta sspB::aadA$<br>$\Delta fur::hyg$ pAB6 | | SAMN12568759 | This study |
| KR4007 | $\Delta lpxC::tetAR \Delta sspB::aadA$ pZIK133 | | | This study |
| KR4008 | $\Delta lpxC::tetAR \Delta sspB::aadA$<br>$\Delta CCNA\_00497::hyg$ pZIK133 | | | This study |
| KR4223 | $\Delta lpxC::tetAR \Delta CCNA\_00497::hyg$<br>pZIK133 | Parent strain for<br>isolating KR4224 and<br>KR4225 | | This study |
| KR4009 | $\Delta lpxC::tetAR \Delta sspB::aadA$<br>$\Delta CCNA\_01553::hyg$ pZIK133 | | | This study |
| KR4010 | $\Delta lpxC::tetAR \Delta sspB::aadA$<br>$\Delta CCNA\_03733::hyg$ pZIK133 | | | This study |
| KR4091 | $\Delta lpxC::tetAR \Delta sspB::aadA$<br>$\Delta fur::hyg$ pZIK133 | | SAMN12568760 | This study |
| KR4113 | $\Delta ctpA::tetAR \Delta sspB::aadA$<br>$\Delta CCNA\_00497::hyg$ | | SAMN12568767 | This study |
| KR4114 | $\Delta ctpA::tetAR \Delta sspB::aadA$<br>$\Delta CCNA\_01553::hyg$ | | SAMN12568768 | This study |
| KR4104 | $\Delta ctpA::tetAR \Delta sspB::aadA$<br>$\Delta CCNA\_03733::hyg$ | | SAMN12568766 | This study |
| KR4102 | $\Delta ctpA::tetAR \Delta sspB::aadA$<br>$\Delta fur::hyg$ | | SAMN12568764 | This study |
| KR4103 | $\Delta lpxC::tetAR \Delta sspB::aadA$<br>$\Delta fur::hyg$ | | SAMN12568765 | This study |
| KR4176 | $\Delta ctpA::tetAR \Delta sspB::aadA$<br>$\Delta fur::hyg$ pZIK175 | | | This study |
| KR4270 | $\Delta ctpA::tetAR \Delta sspB::aadA$<br>$\Delta fur::hyg$ pZIK197 | | | This study |
| KR4177 | $\Delta ctpA::tetAR \Delta sspB::aadA$<br>$\Delta fur::hyg$ pXCERN-2 | | | This study |
| KR4178 | $\Delta lpxC::tetAR \Delta sspB::aadA$<br>$\Delta fur::hyg$ pZIK175 | | | This study |
| KR4271 | $\Delta lpxC::tetAR \Delta sspB::aadA$<br>$\Delta fur::hyg$ pZIK197 | | | This study |
| KR4179 | $\Delta lpxC::tetAR \Delta sspB::aadA$<br>$\Delta fur::hyg$ pXCERN-2 | | | This study |
| KR4180 | $\Delta ctpA::tetAR \Delta sspB::aadA$<br>$\Delta CCNA\_03733::hyg$ pZIK174 | | | This study |
| KR4181 | $\Delta ctpA::tetAR \Delta sspB::aadA$<br>$\Delta CCNA\_03733::hyg$ pXCERN-2 | | | This study |
| KR4182 | $\Delta ctpA::tetAR \Delta sspB::aadA$<br>$\Delta CCNA\_00497::hyg$ pZIK172 | | | This study |
| KR4183 | $\Delta ctpA::tetAR \Delta sspB::aadA$<br>$\Delta CCNA\_00497::hyg$ pXCERN-2 | | | This study |
| KR4148 | $\Delta lpxC::tetAR \Delta sspB::aadA$<br>$\Delta fur::hyg$ pZIK133 | Re-introduction of<br>plasmid for<br>complementation | | This study |
| KR4149 | $\Delta lpxC::tetAR \Delta sspB::aadA$<br>$\Delta fur::hyg$ pJS14 | | | This study |
| KR4150 | $\Delta ctpA::tetAR \Delta sspB::aadA$<br>$\Delta CCNA\_03733::hyg$ pAB6 | Re-introduction of<br>plasmid for<br>complementation | | This study |
| KR4151 | $\Delta ctpA::tetAR \Delta sspB::aadA$<br>$\Delta CCNA\_03733::hyg$ pJS14 | | | This study |
| KR4264 | NA1000 pZIK200 |  |  | This study |
| KR4147 | NA1000 pZIK171 |  |  | This study |
| KR4170 | NA1000 pZIK179 |  |  | This study |
| KR4489 | $\Delta CCNA\_01217$ | | | This study |
| KR4549 | $\Delta CCNA\_01217 vanA::01217::FLAG$ | pKR438 integrated | | This study |
| KR4450 | $\Delta CCNA\_01218$ | | | This study |
| KR4505 | $\Delta CCNA\_01218 vanA::01218::FLAG$ | pKR435 integrated | | This study |
| KR4430 | $\Delta CCNA\_01219$ | | | This study |
| KR4501 | $\Delta CCNA\_01219 vanA::01219::FLAG$ | pKR436 integrated | | This study |
| EK720<br>(KR443<br>1) | $\Delta CCNA\_01220$ | | | (Stankeviciute<br>et al., 2019) |
| KR4530 | $\Delta CCNA\_01220 vanA::01220::FLAG$ | pKR437 integrated | | This study |
| KR4517 | $\Delta fur::hyg \Delta sspB::aadA$<br>$\Delta CCNA\_01217$ | | | This study |
| KR4551 | *Δfur::hyg ΔsspB::aadA*<br>*ΔCCNA_01217 vanA::01217::FLAG* | pKR438 integrated |  | This study |
| KR4442 | *Δfur::hyg ΔsspB::aadA*<br>*ΔCCNA_01218* |  |  | This study |
| KR4503 | *Δfur::hyg ΔsspB::aadA*<br>*ΔCCNA_01218 vanA::01218::FLAG* | pKR435 integrated |  | This study |
| KR4438 | *Δfur::hyg ΔsspB::aadA*<br>*ΔCCNA_01219* |  |  | This study |
| KR4513 | *Δfur::hyg ΔsspB::aadA*<br>*ΔCCNA_01219 vanA::01219::FLAG* | pKR436 integrated |  | This study |
| KR4439 | *Δfur::hyg ΔsspB::aadA*<br>*ΔCCNA_01220* |  |  | This study |
| KR4532 | *Δfur::hyg ΔsspB::aadA*<br>*ΔCCNA_01220 vanA::01220::FLAG* | pKR437 integrated |  | This study |
| KR4509 | *ΔlpxC::tetAR ΔsspB::aadA*<br>*Δfur::hyg ΔCCNA_01217 pZIK133* |  |  | This study |
| KR4555 | *ΔlpxC::tetAR ΔsspB::aadA*<br>*Δfur::hyg ΔCCNA_01217*<br>*vanA::01217::FLAG pZIK133* | pKR438 integrated |  | This study |
| KR4451 | *ΔlpxC::tetAR ΔsspB::aadA*<br>*Δfur::hyg ΔCCNA_01218 pZIK133* |  |  | This study |
| KR4527 | *ΔlpxC::tetAR ΔsspB::aadA*<br>*Δfur::hyg ΔCCNA_01218*<br>*vanA::01218::FLAG pZIK133* | pKR435 integrated |  | This study |
| KR4441 | *ΔlpxC::tetAR ΔsspB::aadA*<br>*Δfur::hyg ΔCCNA_01220 pZIK133* |  |  | This study |
| KR4545 | *ΔlpxC::tetAR ΔsspB::aadA*<br>*Δfur::hyg ΔCCNA_01220*<br>*vanA::01220::FLAG pZIK133* | pKR437 integrated |  | This study |
| KR4205 | *ΔctpA::tetAR ΔsspB::aadA*<br>suppressor isolate #3 | Additional genotype in<br>Supplementary Table 1 | SAMN11107060 | This study |
| KR4206 | *ΔctpA::tetAR ΔsspB::aadA*<br>suppressor isolate #8 | Additional genotype in<br>Supplementary Table 1 | SAMN11107061 | This study |
| KR4207 | *ΔctpA::tetAR ΔsspB::aadA*<br>suppressor isolate #16 | Additional genotype in<br>Supplementary Table 1 | SAMN11107062 | This study |
| KR4208 | *ΔctpA::tetAR ΔsspB::aadA*<br>suppressor isolate #21 | Additional genotype in<br>Supplementary Table 1 | SAMN11107063 | This study |
| KR4209 | *ΔctpA::tetAR ΔsspB::aadA*<br>suppressor isolate #32 | Additional genotype in<br>Supplementary Table 1 | SAMN11107064 | This study |
| KR4210 | *ΔctpA::tetAR ΔsspB::aadA*<br>suppressor isolate #36 | Additional genotype in<br>Supplementary Table 1 | SAMN11107065 | This study |
| KR4211 | *ΔctpA::tetAR ΔsspB::aadA*<br>suppressor isolate #38 | Additional genotype in<br>Supplementary Table 1 | SAMN11107066 | This study |
| KR4212 | *ΔctpA::tetAR ΔsspB::aadA*<br>suppressor isolate #40 | Additional genotype in<br>Supplementary Table 1 | SAMN11107067 | This study |
| KR4213 | *ΔctpA::tetAR ΔsspB::aadA*<br>suppressor isolate #43 | Additional genotype in<br>Supplementary Table 1 | SAMN11107068 | This study |
| KR4214 | $\Delta ctpA::tetAR \Delta sspB::aadA$ suppressor isolate #44 | Additional genotype in Supplementary Table 1 | SAMN11107069 | This study |
| KR4215 | $\Delta ctpA::tetAR \Delta sspB::aadA$ suppressor isolate #47 | Additional genotype in Supplementary Table 1 | SAMN11107070 | This study |
| KR4216 | $\Delta ctpA::tetAR \Delta sspB::aadA$ suppressor isolate #52 | Additional genotype in Supplementary Table 1 | SAMN11107071 | This study |
| KR4217 | $\Delta ctpA::tetAR \Delta sspB::aadA$ suppressor isolate #53 | Additional genotype in Supplementary Table 1 | SAMN11107072 | This study |
| KR4218 | $\Delta ctpA::tetAR \Delta sspB::aadA$ suppressor isolate #54 | Additional genotype in Supplementary Table 1 | SAMN11107073 | This study |
| KR4219 | $\Delta ctpA::tetAR \Delta sspB::aadA$ suppressor isolate #57 | Additional genotype in Supplementary Table 1 | SAMN11107074 | This study |
| KR4220 | $\Delta ctpA::tetAR \Delta sspB::aadA$ suppressor isolate #111 | Additional genotype in Supplementary Table 1 | SAMN11107075 | This study |
| KR4221 | $\Delta ctpA::tetAR \Delta sspB::aadA$ suppressor isolate #112 | Additional genotype in Supplementary Table 1 | SAMN11107076 | This study |
| KR4224 | $\Delta lpxC::tetAR \Delta sspB::aadA$ suppressor isolate #1 | Additional genotype in Supplementary Table 2 | SAMN11107077 | This study |
| KR4225 | $\Delta lpxC::tetAR \Delta sspB::aadA$ suppressor isolate #5 | Additional genotype in Supplementary Table 2 | SAMN11107078 | This study |

**Table S5, related to STAR Methods:**
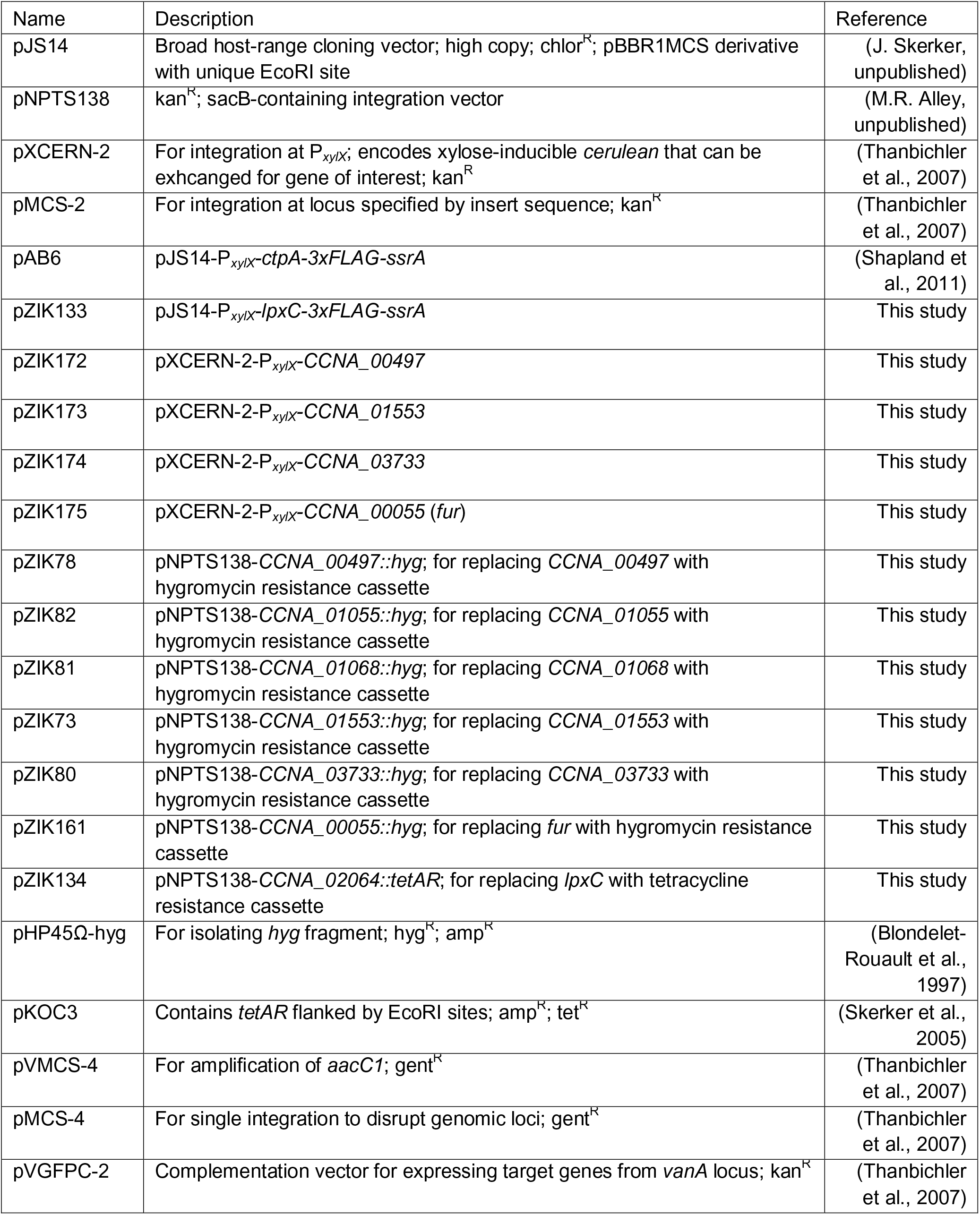
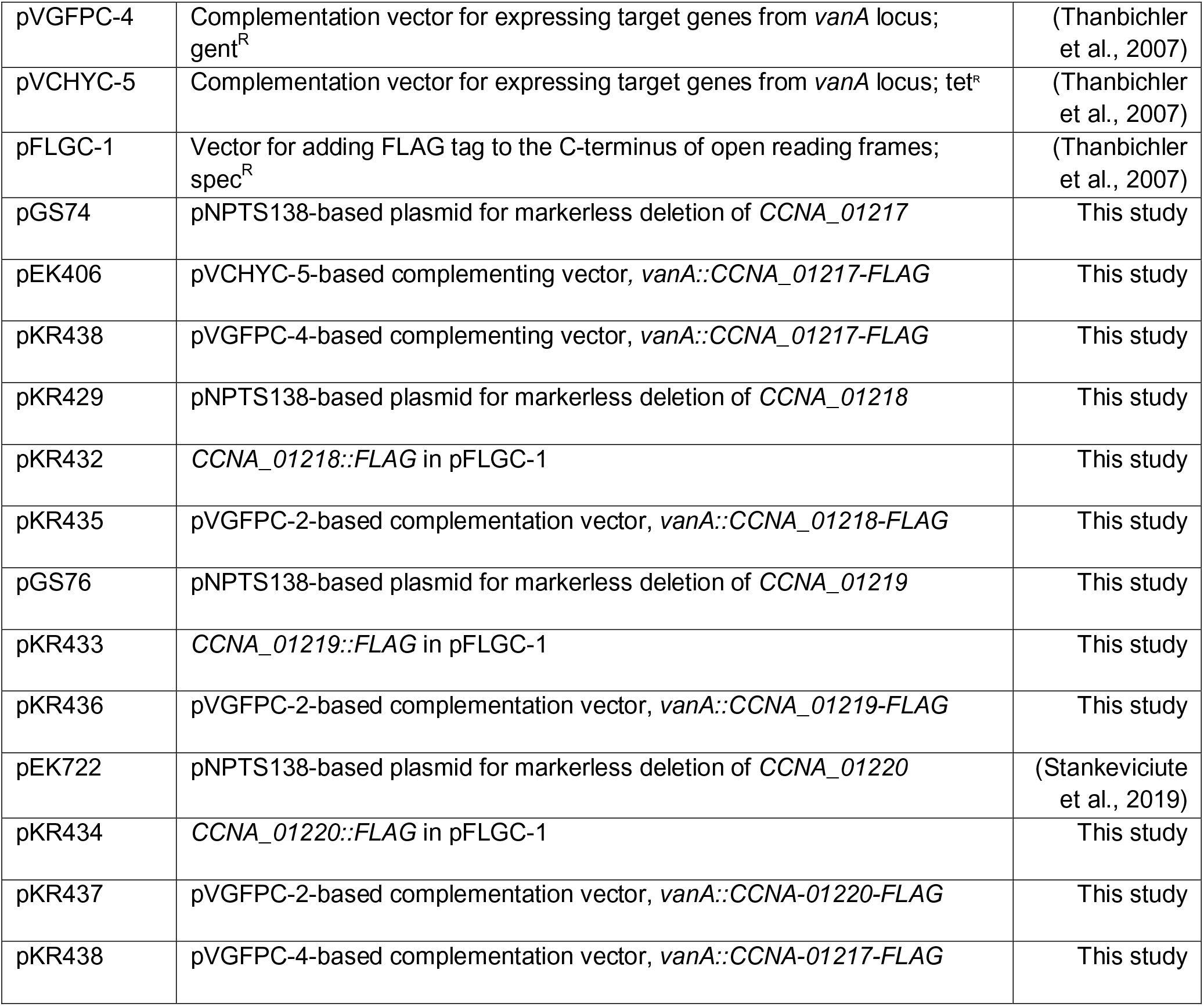
Plasmids used in this study.

**Table S6, related to STAR Methods:**
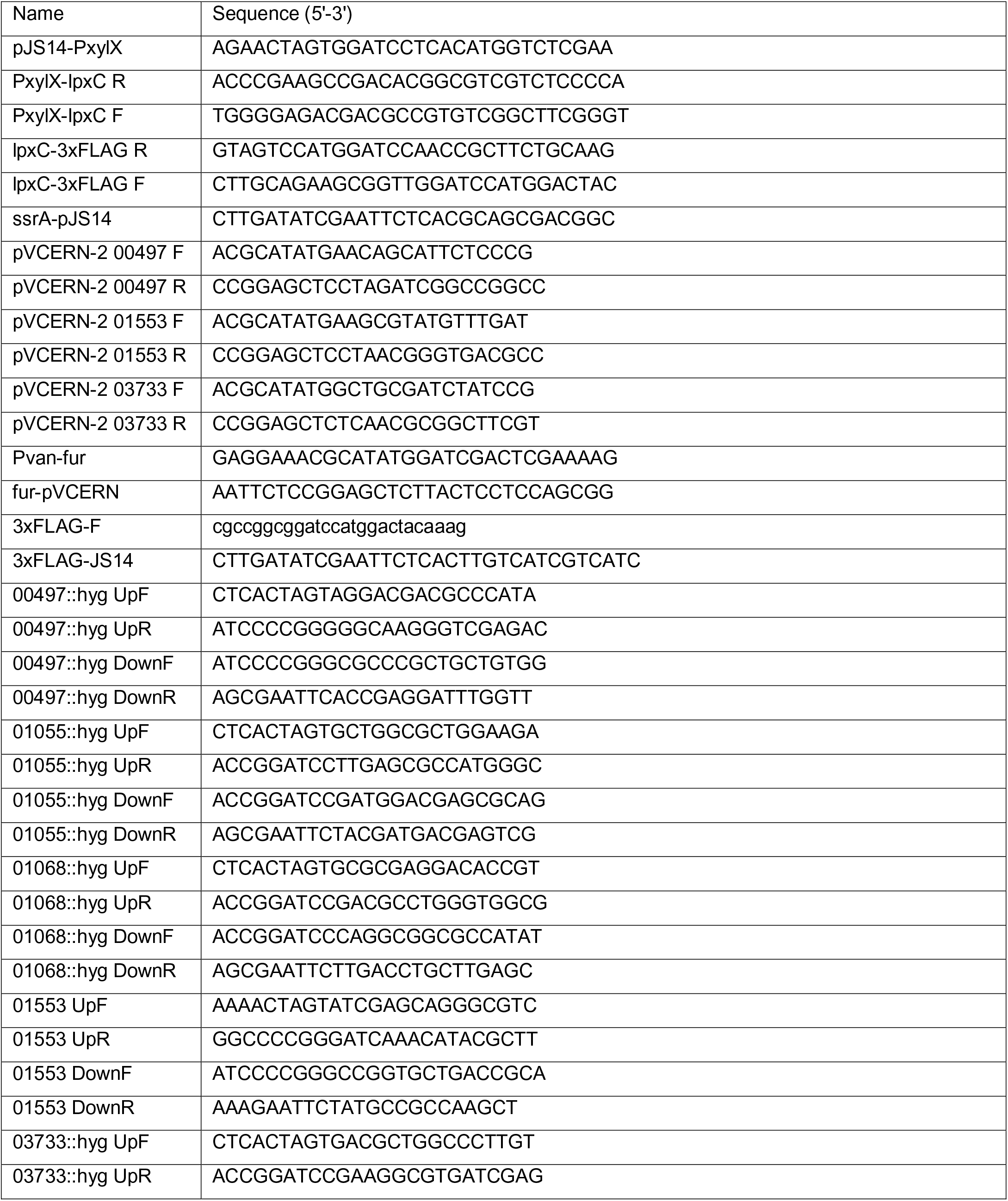
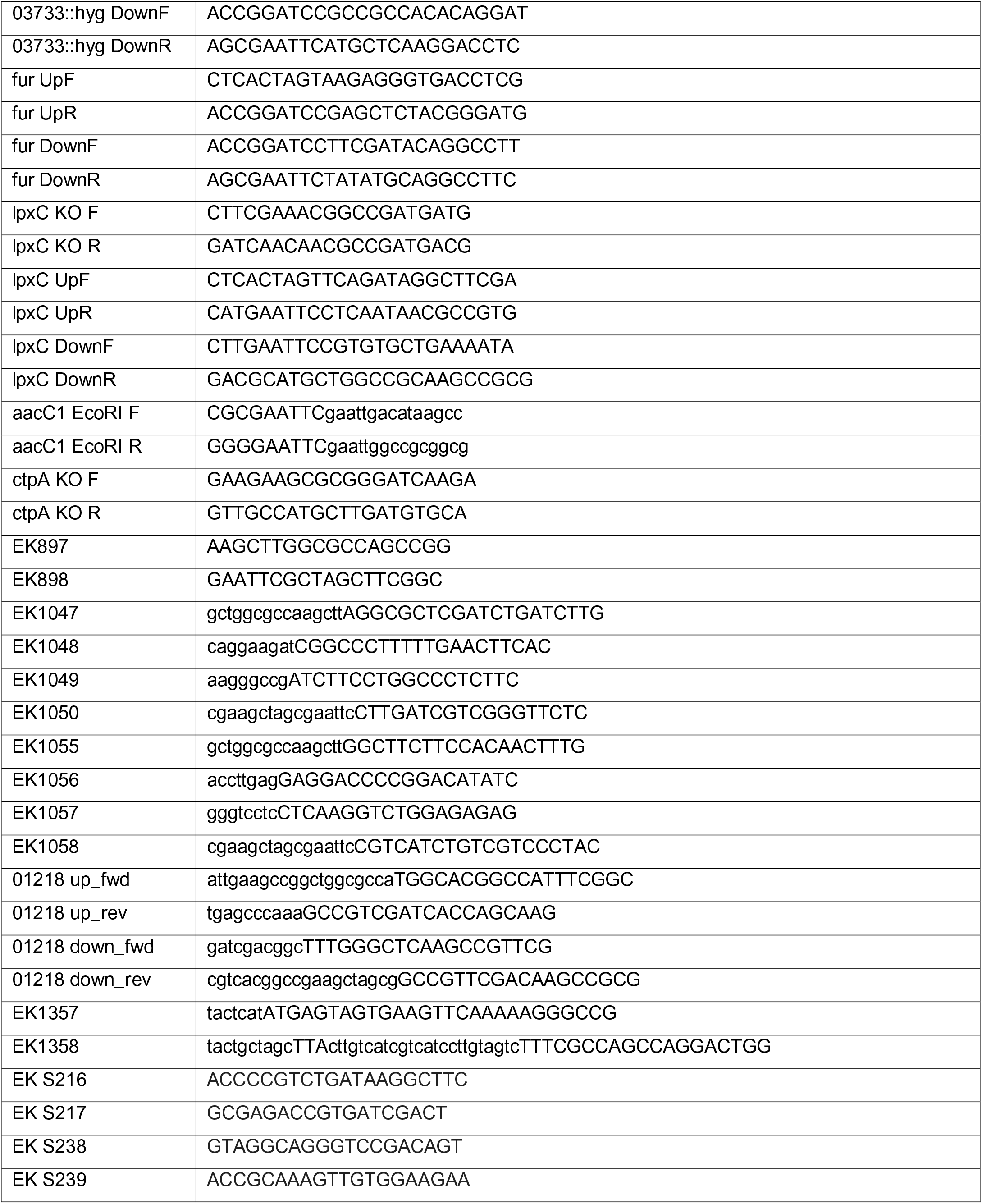
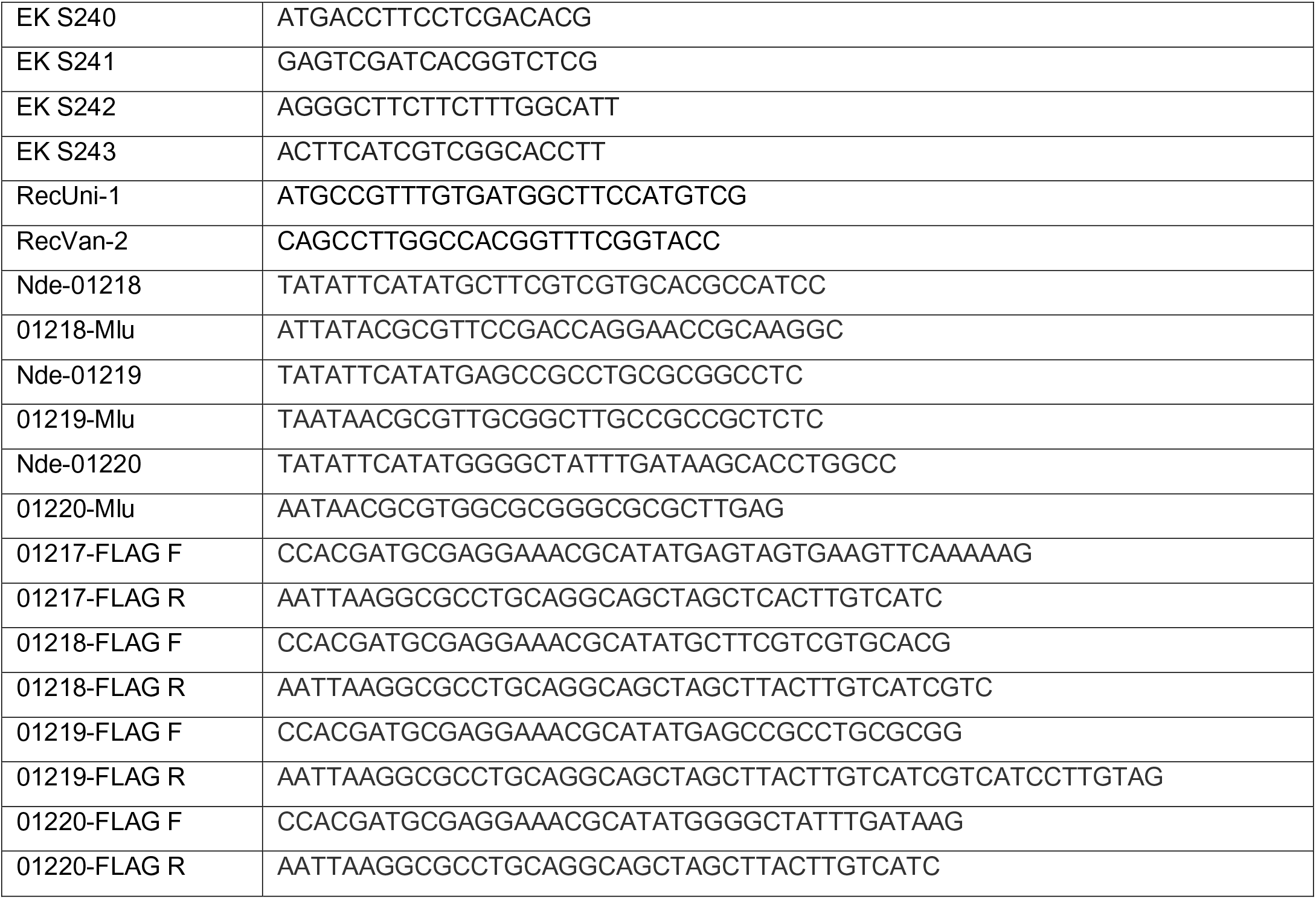
Primers used in this study.

## References

Afgan, E., Baker, D., Beek, M. van den, Blankenberg, D., Bouvier, D., Cech, M., Chilton, J., Clements, D., Coraor, N., Eberhard, C., et al. (2016). The Galaxy platform for accessible, reproducible and collaborative biomedical analyses: 2016 update. Nucleic Acids Res. 44, W3– W10.

Andrews, S., Norton, I., Salunkhe, A.S., Goodluck, H., Aly, W.S.M., Mourad-Agha, H., and Cornelis, P. (2013). Control of iron metabolism in bacteria. Met. Ions Life Sci. 12, 203–239.

Awram, P., and Smit, J. (2001). Identification of lipopolysaccharide O antigen synthesis genes required for attachment of the S-layer of Caulobacter crescentus. Microbiology 147, 1451–1460.

Bligh, E.G., and Dyer, W.J. (1959). A rapid method of total lipid extraction and purification. Can J. Biochem. Physiol. 37, 911–917.

Blondelet-Rouault, M.H., Weiser, J., Lebrihi, A., Branny, P., and Pernodet, J.L. (1997). Antibiotic resistance gene cassettes derived from the omega interposon for use in E. coli and Streptomyces. Gene 190, 315–317.

Boll, J.M., Crofts, A.A., Peters, K., Cattior, V., Vollmer, W., Davies, B.W., and Trent, M.S. (2016). A penicillin-binding protein inhibits selection of colistin-resistant, lipooligosaccharide-deficient Acinetobacter baumannii. Proc. Natl. Acad. Sci. USA 113, E6228–E6237.

Brown, D.B., Forsberg, L.S., Kannenberg, E.L., and Carlson, R.W. (2012). Characterization of galacturonosyl transferase genes rgtA, rgtB, rgtC, rgtD, and rgtE responsible for lipopolysaccharide synthesis in nitrogen-fixing endosymbiont Rhizobium leguminosarum: lipopolysaccharide core and lipid galacturonosyl residues confer membrane stability. J. Biol. Chem. 287, 935–949.

Brown, D.B., Muszynski, A., and Carlson, R.W. (2013). Elucidation of a novel lipid A alpha-(1,1)-GalA transferase gene (rgtF) from Mesorhizobium loti: Heterologous expression of rgtF causes Rhizobium etli to synthesize lipid A with alpha-(1,1)-GalA. Glycobiology 23, 546–558.

Christen, B., Abeliuk, E., Collier, J.M., Kalogeraki, V.S., Passarelli, B., Coller, J.A., Fero, M.J., McAdams, H.H., and Shapiro, L. (2011). The essential genome of a bacterium. Mol. Syst. Biol. 7, 528.

Crosson, S., McGrath, P.T., Stephens, C., McAdams, H.H., and Shapiro, L. (2005). Conserved modular design of an oxygen sensory/signaling network with species-specific output. Proc. Natl. Acad. Sci. USA 102, 8018–8023.

Darveau, R.P., and Hancock, R.E. (1983). Procedure for isolation of bacterial lipopolysaccharides from both smooth and rough Pseudomonas aeruginosa and Salmonella typhimurium strains. J. Bacteriol. 155, 831–838.

Davis, M.R.J., and Goldberg, J.B. (2012). Purification and visualization of lipopolysaccharide from Gram-negative bacteria by hot aqueous-phenol extraction. J. Vis. Exp. e3916.

De Castro, C., Molinaro, A., Lanzetta, R., Silipo, A., and Parrilli, M. (2008). Lipopolysaccharide structures from Agrobacterium and Rhizobiaceae species. Carbohydr. Res. 343, 1924–1933.

Dehio, C., and Meyer, M. (1997). Maintenance of broad-host-range incompatibility group P and group Q plasmids and transposition of Tn5 in Bartonella henselae following conjugal plasmid transfer from Escherichia coli. J. Bacteriol. 179, 538–540.

El Hamidi, A., Tirsoaga, A., Novikov, A., Hussein, A., and Caroff, M. (2005). Microextraction of bacterial lipid A: Easy and rapid method for mass spectrometric characterization. J. Lipid Res. 46, 1773–1778.

Ely, B. (1991). Genetics of Caulobacter crescentus. Methods Enzymol. 204, 372–384.

Evinger, M., and Agabian, N. (1977). Envelope-associated nucleoid from Caulobacter crescentus stalked and swarmer cells. J. Bacteriol. 132, 294–301.

Fontenot, C.R., Tasnim, H., Valdes, K.A., Popescu, C.V., and Ding, H. (2020). Ferric uptake regulator (Fur) reversibly binds a [2Fe-2S] cluster to sense intracellular iron homeostasis in Escherichia coli. J. Biol. Chem. 295, 15454–15463.

Frangakis, A.S., and Hegerl, R. (2001). Noise reduction in electron tomographic reconstructions using nonlinear anisotropic diffusion. J. Struct. Biol. 135, 239–250.

Gilchrist, A., and Smit, J. (1991). Transformation of freshwater and marine caulobacters by electroporation. J. Bacteriol. 173, 921–925.

Guan, Z., Katzianer, D., Zhu, J., and Goldfine, H. (2014). Clostridium difficile contains plasmalogen species of phospholipids and glycolipids. Biochim. Biophys. Acta 1842, 1353– 1359.

Hassett, D.J., Britigan, B.E., Svendsen, T., Rosen, G.M., and Cohen, M.S. (1987). Bacteria form intracellular free radicals in response to paraquat and streptonigrin. Demonstration of the potency of hydroxyl radical. J. Biol. Chem. 262, 13404–13408.

Henry, R., Crane, B., Powell, D., Lucas, D.D., Li, Z., Aranda, J., Harrison, P., Nation, R.L., Adler, B., Harper, M., et al. (2015). The transcriptomic response of Acinetobacter baumannii to colistin and diripenem alone and in combination in an in vitro pharmacokinetics/pharmacodynamics model. J. Antimicrob. Chermother. 70, 1303–1313.

Hershey, D.M., Fiebig, A., and Crosson, S. (2019). A genome-wide analysis of adhesion in Caulobacter crescentus identifies new regulatory and biosynthetic components for holdfast assembly. mBio 10, 2273.

Jones, M.D., Vinogradov, E., Nomellini, J.F., and Smit, J. (2015). The core and O-polysaccharide structure of the Caulobacter crescentus lipopolysaccharide. Carbohydr. Res. 402, 111–117.

Justino, M.C., Almeida, C.C., Teixeira, M., and Saraiva, L.M. (2007). Escherichia coli di-iron YtfE protein is necessary for the repair of stress-damaged iron-sulfur clusters. J. Biol. Chem. 282, 10352–10359.

Karbarz, M.J., Kalb, S.R., Cotter, R.J., and Raetz, C.R.H. (2003). Expression cloning and biochemical characterization of a Rhizobium leguminosarum lipid A 1-phosphatase. J. Biol. Chem 278, 39269–39279.

Kawahara, K., Seydel, U., Matsuura, M., Danbara, H., Rietschel, E.T., and Zahringer, U. (1991). Chemical structure of glycosphingolipids isolated from Sphingomonas paucimobilis. FEBS Lett. 292, 107–110.

Kawasaki, S., Moriguchi, R., Sekiya, K., Nakai, T., Ono, E., Kume, K., and Kawahara, K. (1994). The cell envelope structure of the lipopolysaccharide-lacking Gram-negative bacterium Sphingomonas paucimobilis. J. Bacteriol. 176, 284–290.

Kremer, J.R., Mastonarde, D.N., and McIntosh, J.R. (1996). Computer visualization of three-dimensional image data using IMOD. J. Struct. Biol. 116, 71–76.

Leaden, L., Silva, L.G., Ribiero, R.A., Santos, N.M.D., Lorenzetti, A.P.R., Alegria, T.G.P., Schultz, M.L., Medeiros, M.H.G., Koide, T., and Marques, M.V. (2018). Iron deficiency generates oxidative stress and activation of the SOS response in Caulobacter crescentus. Front. Microbiol. 9, 2014.

Leung, L.M., Fondrie, W.E., Doi, Y., Johnson, J.K., Strickland, D.K., Ernst, R.K., and Goodlett, D.R. (2017). Identification of the ESKAPE pathogens by mass spectrometric analysis of microbial membrane glycolypids. Sci. Rep. 7, 6403.

Levchenko, I., Seidel, M., Sauer, R.T., and Baker, T.A. (2000). A specificity-enhancing factor for the ClpXP degradation machine. Science 289, 2354–2356.

Liu, L., Feng, X., Wang, W., Chen, Y., Chen, Z., and Gao, H. (2020). Free Rather Than Total Iron Content Is Critically Linked to the Fur Physiology in Shewanella oneidensis. Front. Microbiol. 11, 593246.

Mamat, U., Meredith, T.C., Aggarwal, P., Kuhl, A., Kirchoff, P., Lindner, B., Hanuszkiewicz, A., Sun, J., Holst, O., and Woodard, R.W. (2008). Single amino acid substitutions in either YhjD or MsbA confer viability to 3-deoxy-D-manno-oct-2-ulosonic acid-depleted Escherichia coli. Mol. Microbiol. 67, 633–648.

Marks, M.E., Castro-Rojas, C., Teiling, C., Du, L., Kapatral, V., Walunas, T.L., and Crosson, S. (2010). The genetic basis of laboratory adaptation in Caulobacter crescentus. J. Bacteriol. 192, 3678–3688.

Mastonarde, D.N. (2005). Automated electron microscope tomography using robust prediction of specimen movements. J. Struct. Biol. 152, 36–51.

McClerren, A.L., Endsley, S., Bowman, J.L., Andersen, N.H., Guan, Z., Rudolph, J., and Raetz, C.R. (2005). A slow, tight-binding inhibitor of the zinc-dependent deacetylase LpxC of lipid A biosynthesis with antibiotic activity comparable to ciprofloxacin. Biochemistry 44, 16574–16583.

Meisenzahl, A.C., Shapiro, L., and Jenal, U. (1997). Isolation and characterization of a xylose-dependent promoter from Caulobacter crescentus. J. Bacteriol. 179, 592–600.

Meredith, T.C., Aggarwal, P., Mamat, U., Lindner, B., and Woodard, R.W. (2006). Redefining the requisite lipopolysaccharide structure in Escherichia coli. ACS Chem. Biol. 1, 33–42.

Moffatt, J.H., Harper, M., Harrison, P., Hale, J.D., Vinogradov, E., Seemann, T., Henry, R., Crane, B., Michael, F.S., Cox, A.D., et al. (2010). Colistin resistance in Acinetobacter baumannii is mediated by complete loss of lipopolysaccharide production. Antimicrob. Agents Chemother. 54, 4971–4977.

Moffatt, J.H., Harper, M., and Boyce, J.D. (2019). Mechanisms of polymyxin resistance. Adv. Exp. Med. Biol. 55–71.

Nachin, L., El Hassouni, M., Loiseau, L., Expert, D., and Barras, F. (2001). SoxR-dependent response to oxidative stress and virulence of Erwinia chrysanthemi: the key role of SufC, an orphan ABC ATPase. Mol. Microbiol. 39, 960–972.

Nagiec, M.M., Skrzypek, M., Nagiec, E.E., Lester, R.L., and Dickson, R.C. (1998). The LCB4 (YOR171c) and LCB5 (YLR260w) genes of Saccharomyces encode sphingoid long chain base kinases. J. Biol. Chem. 273, 19437–19442.

Nagy, E., Losick, R., and Kahne, D. (2019). Robust suppression of lipopolysaccharide deficiency in Acinetobacter baumannii by growth in minimal medium. J. Bacteriol. 201, 420.

Nikaido, H. (2003). Molecular basis of bacterial outer membrane permeability revisited. Microbiol. Mol. Biol. Rev. 67, 593–656.

Olea-Ozuna, R.J., Poggio, S., Bergström, E., Quiroz-Rocha, E., García-Soriano, D.A., Sahonero-Canavesi, D.X., Padilla-Gómez, J., Martínez-Aguilar, L., López-Lara, I.M., Thomas-Oates, J., et al. (2021). Five structural genes required for ceramide synthesis in Caulobacter and for bacterial survival. Environ. Microbiol. 23, 143–159.

Peng, D., Hong, W., Choudhury, B.P., Carlson, R.W., and Gu, X.-X. (2005). Moraxella cararrhalis bacterium without endotoxin, a potential vaccine candidate. Infect. Immun. 73, 7569– 7577.

Plötz, B.M., Lindner, B., Stetter, K.O., and Holst, O. (2000). Characterization of a novel lipid A containing D-galacturonic acid that replaces phosphate residues. The structure of the lipid a of the lipopolysaccharide from the hyperthermophilic bacterium Aquifex pyrophilus. J. Biol. Chem. 275, 11222–11228.

Price, M.N., Wetmore, K.M., Waters, R.J., Callaghan, M., Ray, J., Liu, H., Kuehl, J.V., Melnyk, R.A., Lamson, J.S., Suh, Y., et al. (2018). Mutant phenotypes for thousands of bacterial genes of unknown function. Nature 557, 503–509.

Qureshi, N., Takayama, K., Mascagni, P., Honovich, J., Wong, R., and Cotter, R.J. (1988). Complete structural determination of lipopolysaccharide obtained from deep rough mutant of Escherichia coli. Purification by high performance liquid chromatography and direct analysis by plasma desorption mass spectrometry. J. Biol. Chem. 263, 11971–11976.

Radolf, J.D., and Kumar, S. (2018). The Treponema pallidum outer membrane. Curr. Top. Microbiol. Immuno. 415, 1–38.

Rojas, E.R., Billings, G., Odermatt, P.D., Auer, g K., Zhu, L., Miguel, A., Chang, F., Weibel, D.B., Theriot, J.A., and Huang, K.C. (2018). The outer membrane is an essential load-bearing element in Gram-negative bacteria. Nature 559, 617–621.

Samuel, G., and Reeves, P. (2003). Biosynthesis of O-antigens: Genes and pathways involved in nucleotide sugar precursor synthesis and O-antigen assembly. Carboyhdr. Res. 338, 2503– 2519.

Shapland, E.B., Reisinger, S.J., Bajwa, A.K., and Ryan, K. (2011). An essential tyrosine phosphatase homolog regulates cell separation, outer membrane integrity, and morphology in Caulobacter crescentus. J. Bacteriol. 193, 4361–4370.

da Silva Neto, J.F., Braz, V.S., Italiani, V.C.S., and Marques, M.V. (2009). Fur controls iron homeostasis and oxidative stress defense in the oligotrophic alpha-proteobacterium Caulobacter crescentus. Nucleic Acids Res. 37, 4812–4825.

da Silva Neto, J.F., Lourenco, R.F., and Marques, M.V. (2013). Global transcriptional response of Caulobacter crescentus to iron availability. BMC Genomics 14, 549.

Simpson, B.W., Nieckarz, M., Pinedo, V., McLean, A.B., Cava, F., and Trent, M.S. (2021). Acinetobacter baumannii Can Survive with an Outer Membrane Lacking Lipooligosaccharide Due to Structural Support from Elongasome Peptidoglycan Synthesis. mBio 12, e0309921.

Skerker, J.M., Prasol, M.S., Perchuk, B.S., Biondi, E.G., and Laub, M.T. (2005). Two-component signal transduction pathways regulating growth and cell cycle progresion in a bacterium: A systems-level analysis. PLoS Biol. 3, e334.

Smit, J., Kaltashov, I.A., Cotter, R.J., Vinogradov, E., Perry, M.B., Haider, H., and Qureshi, N. (2008). Structure of a novel lipid A obtained from the lipopolysaccharide of Caulobacter crescentus. Innate Immun. 14, 25–37.

Stankeviciute, G., Guan, Z., Goldfine, H., and Klein, E.A. (2019). Caulobacter crescentus adapts to phosphate starvation by synthesizing anionic glycoglycerolipids and a novel glycosphingolipid. mBio 10, 107.

Stankeviciute, G., Tang, P., Ashley, B., Chamberlain, J.D., Hansen, M.E.B., Coleman, A., D’Emilia, R., Fu, L., Mohan, E.C., Nguyen, H., et al. (2021). Convergent evolution of bacterial ceramide synthesis. Nat. Chem. Biol. DOI: 10.1038/s41589-021-00948-7.

Steeghs, L., Hartog, R. den, Boer, A. den, Zomer, B., Roholl, P., and Ley, P. van der (1998). Meningitis bacterium is viable without endotoxin. Nature 392, 449–450.

Thanbichler, M., Iniesta, A.A., and Shapiro, L. (2007). A comprehensive set of plasmids for vanillate- and xylose-inducible gene expression in Caulobacter crescentus. Nucleic Acids Res. 35, e137.

Toh, E., Jr, H.D.K., and Brun, Y.V. (2008). Characterization of the Caulobacter crescentus holdfast polysaccharide biosynthesis pathway reveals significant redundancy in the initiating glycosyltransferase and polyerase steps. J. Bacteriol. 190, 7219–7131.

Velkov, T., Thompson, P.E., Nation, R.L., and Li, J. (2010). Structure--activity relationships of polymyxin antibiotics. J. Med. Chem. 53, 1898–1916.

Walker, S.G., Karunaratne, D.N., Ravenscroft, N., and Smit, J. (1994). Characterization of mutants of Caulobacter crescentus defective in surface attachment of the paracrystalline surface layer. J. Bacteriol. 176, 6313–6323.

Wang, X., Ribeiro, A.A., Guan, Z., McGrath, S.C., Cotter, R.J., and Raetz, C.R.H. (2006). Structure and biosynthesis of free lipid A molecules that replace lipopolysaccharide in Francisella tularensis subsp. novicida. Biochemistry 45, 14427–14440.

Westphal, O., and Jann, K. (1965). Bacterial lipopolysaccharides extraction with phenol-water and further applications of the procedure. Methods Carbohydr. Chem. 5, 83–91.

Wetmore, K.M., Price, M.N., Waters, R.J., Lamson, J.S., He, J., Hoover, C.A., Blow, M.J., Bristow, J., Butland, G., Arkin, A.P., et al. (2015). Rapid quantification of mutant fitness in diverse bacteria by sequencing randomly bar-coded transposons. mBio 6, e00306–e00315.

Whitfield, C., and Trent, M.S. (2014). Biosynthesis and export of bacterial lipopolysaccharides. Annu. Rev. Biochem. 83, 99–128.

Wofford, J.D., Bolaji, N., Dziuba, N., Outten, F.W., and Lindahl, P.A. (2019). Evidence that a respiratory shield in Escherichia coli protects a low-molecular-mass FeII pool from O2-dependent oxidation. J. Biol. Chem. 294, 50–62.

Yeowell, H.N., and White, J.R. (1982). Iron requirement in the bactericidal mechanism of streptonigrin. Antimicrob. Agents Chemother. 22, 961–968.

Yoon, S.H., Liang, T., Schneider, T., Oyler, B.L., Chandler, C.E., Ernst, R.K., Yen, G.S., Huang, Y., Nilsson, E., and Goodlett, D.R. (2016). Rapid lipid a structure determination via surface acoustic wave nebulization and hierarchical tandem mass spectrometry algorithm. Rapid Commun. Mass Spectrom. 30, 2555–2560.

Zhang, G., Meredith, T.C., and Kahne, D. (2013). On the essentiality of lipopolysaccharide to Gram-negative bacteria. Curr. Opin. Microbiol. 16, 779–785.

Zheng, S.Q., Plaovcak, E., Armache, J.P., Verba, K.A., Cheng, Y., and Agard, D.A. (2017). MotionCor2: anisotropic correction of bean-induced motion for improved cryo-electron microscopy. Nat. Methods 14, 331–332.

Zhou, B., Schrader, J.M., Kalogeraki, V.S., Abeliuk, E., Dinh, C.B., Pham, J.Q., Cui, Z.Z., Dill, D.L., McAdams, H.H., and Shapiro, L. (2015). The global regulatory architecture of transcription during the Caulobacter cell cycle. PLoS Genet. 11, e1004831.

