## Supplementary figures and images for "*Caulobacter* requires anionic sphingolipids and deactivation of *fur* to lose lipid A"

### Supplemental Figure S1

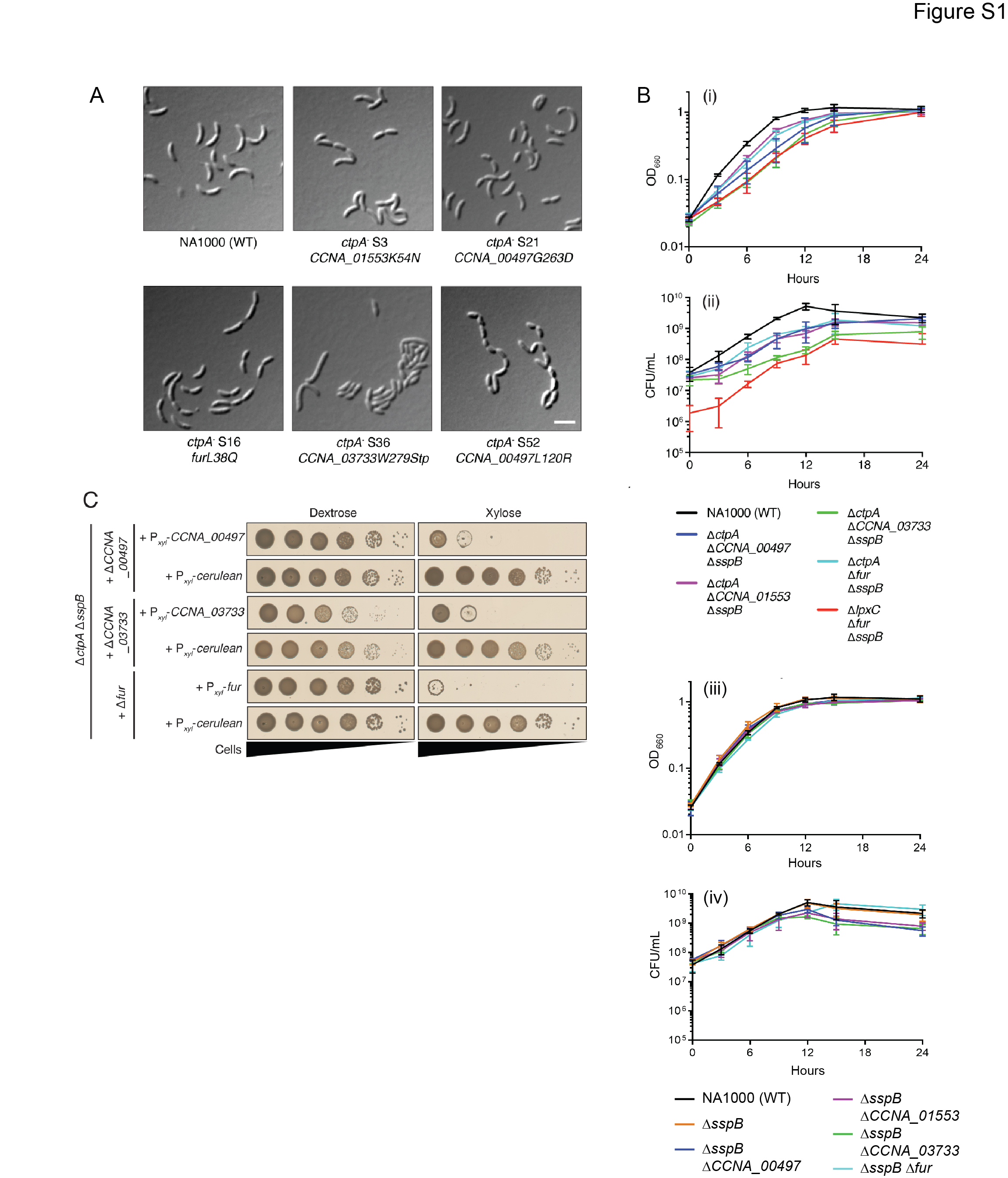

### Supplemental Figure S2

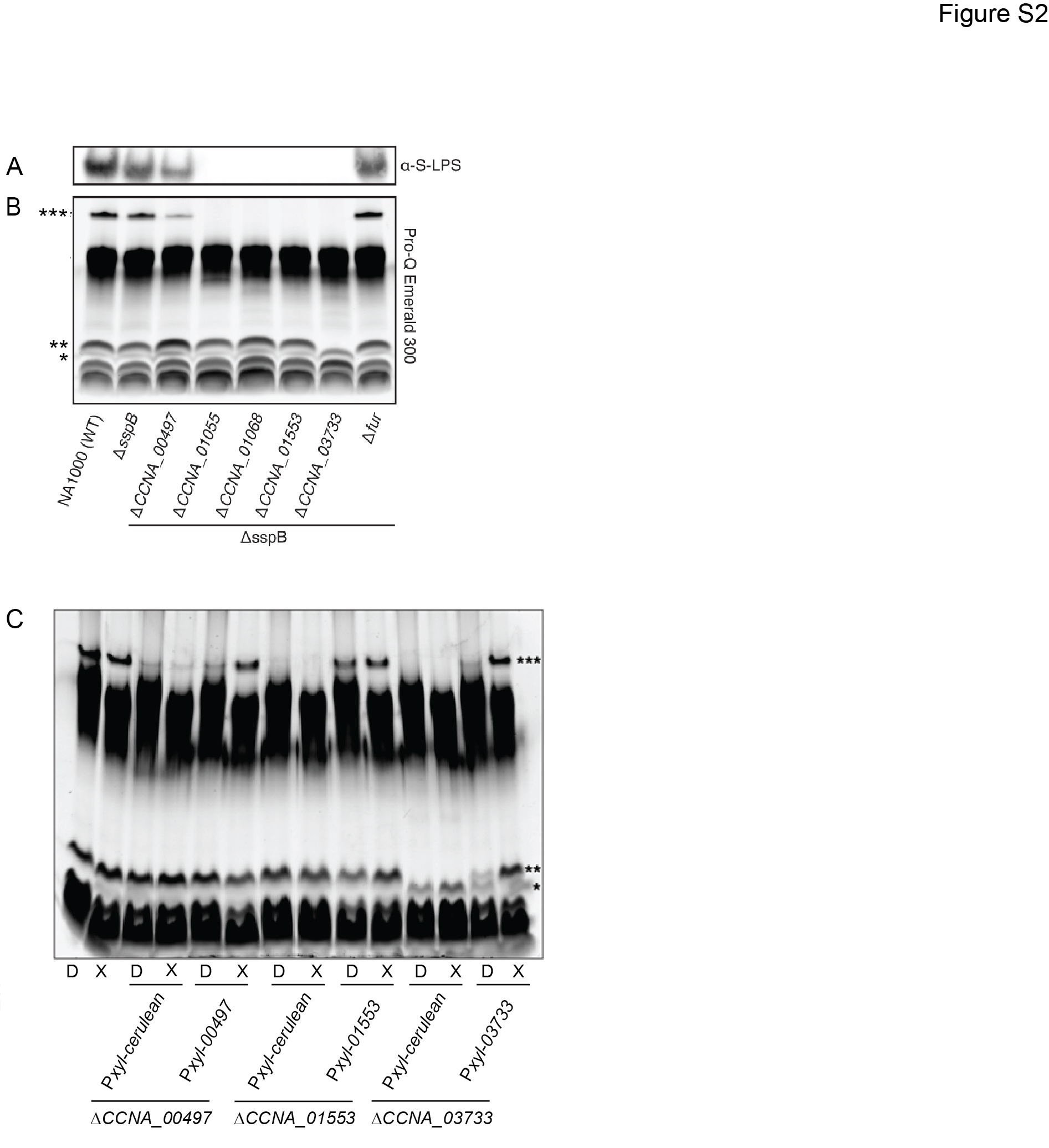

### Supplemental Figure S3

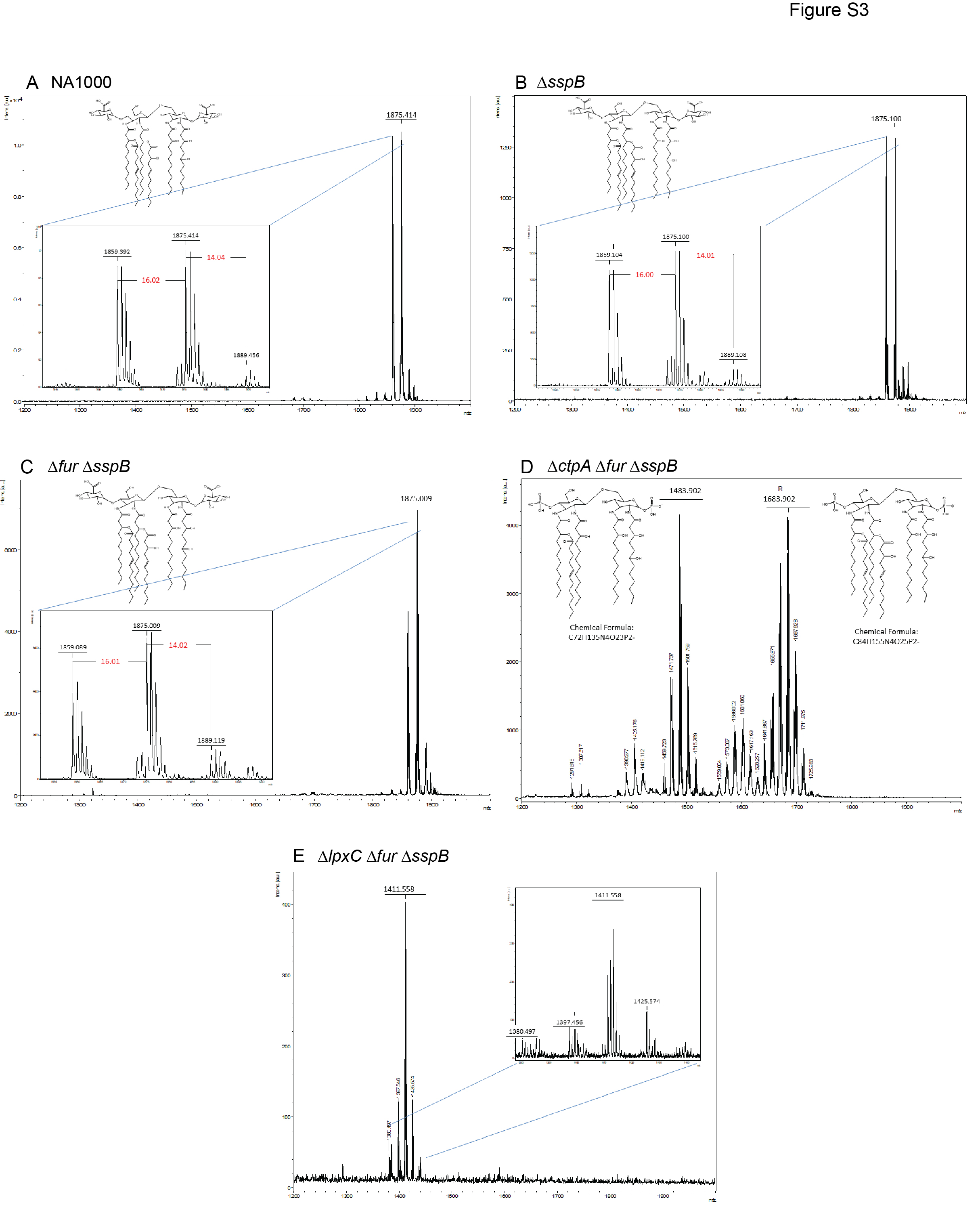

### Supplemental Figure S4

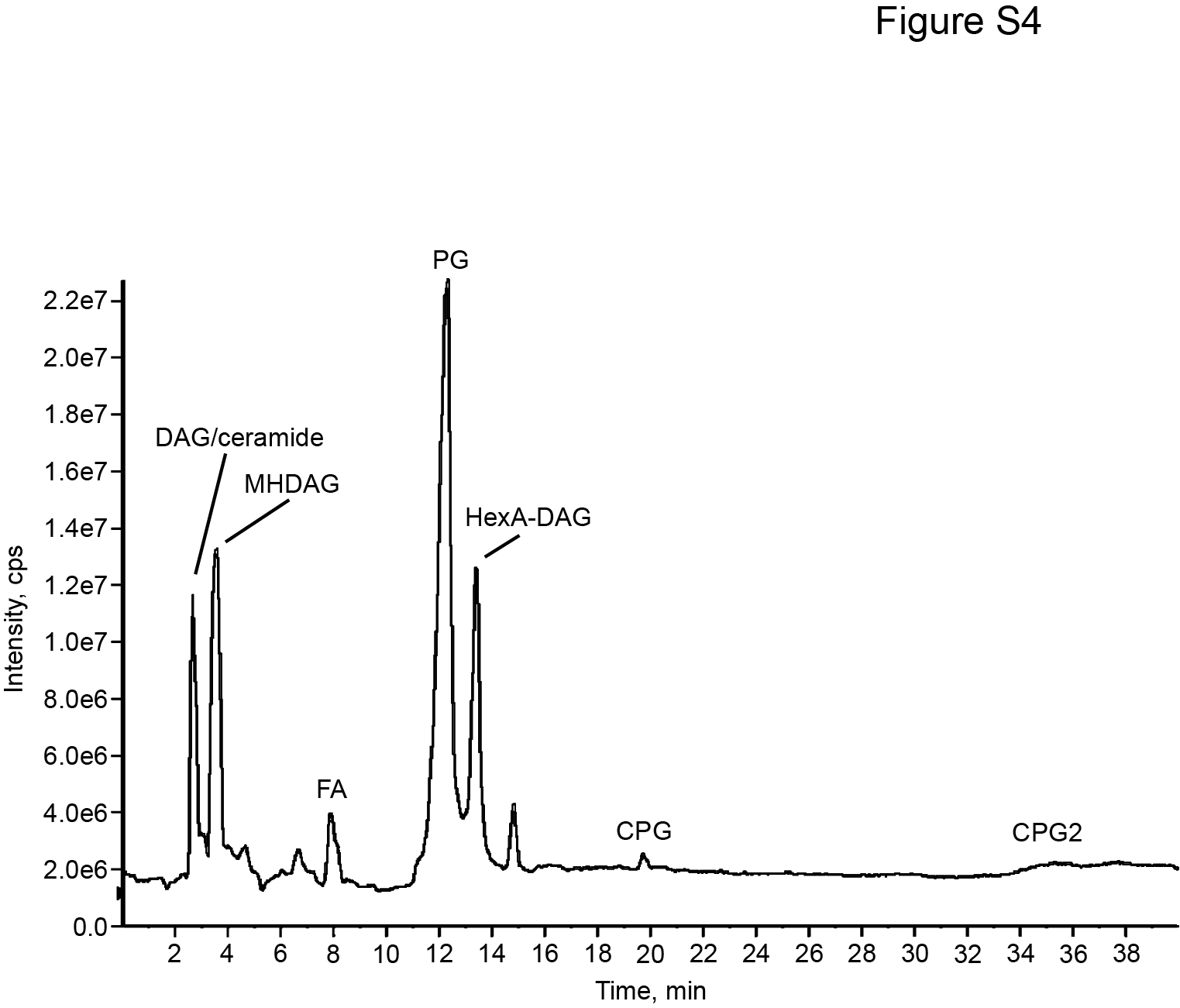

### Supplemental Figure S5

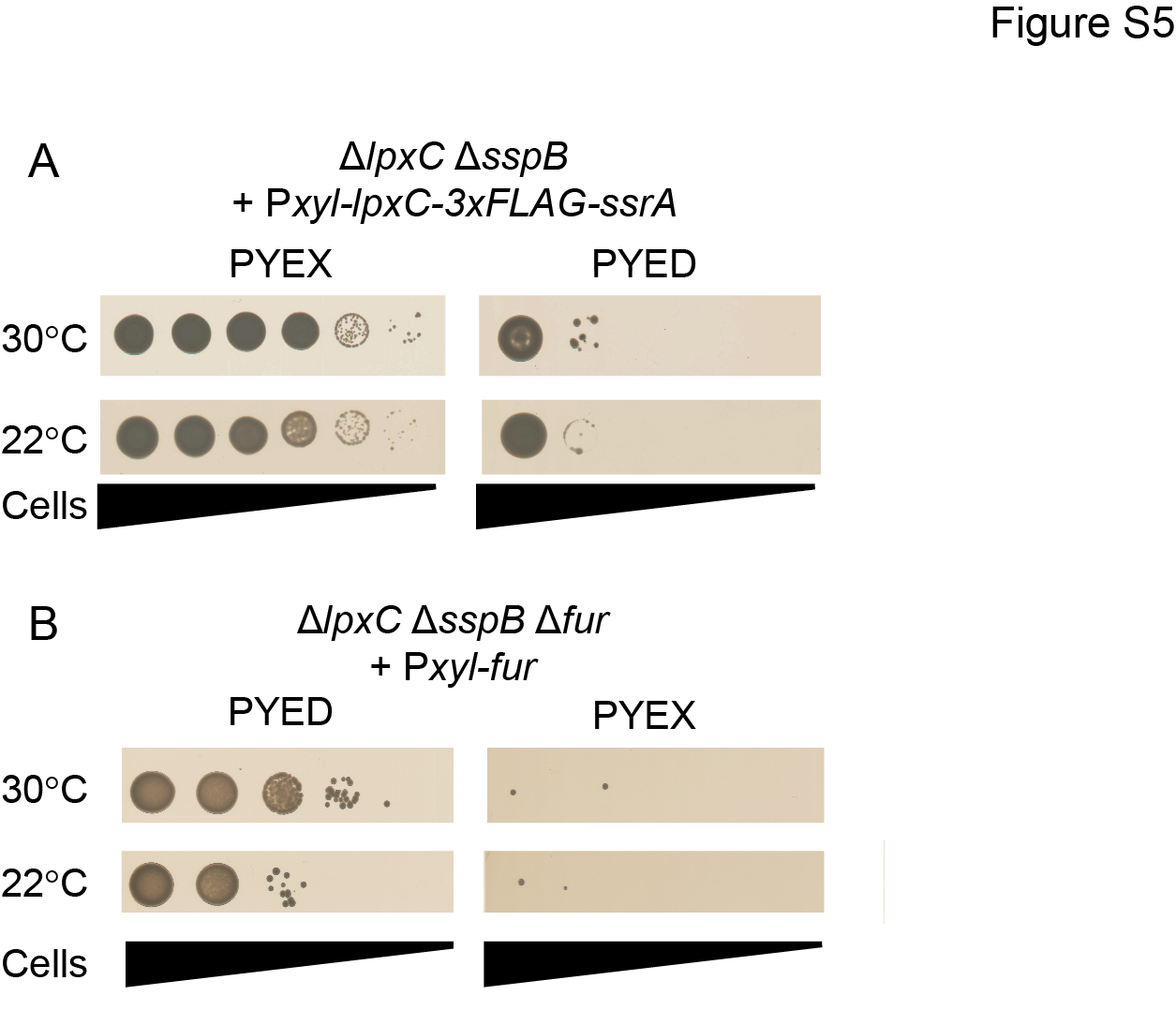
